# A novel allele of *Sh1* underlies the conversion of *waxy* corn to *wx-sweet* corn

**DOI:** 10.1101/2025.11.05.686684

**Authors:** Ling Zhou, Lihua Ning, Shuaiqiang Liang, Qiaoqiang Hu, Wenxiang Zhang, Yuxin Pan, Zhifang Gao, Jun Hong, Yuancong Wang, Wenming Zhao, Yanping Chen, Huixue Dai, Han Zhao

## Abstract

Waxy corn and sweet corn are two major types of fresh-eating corn. Each has significant market value due to its unique sensory and nutritional profiles. Developing a variety that combines both traits would meet growing consumer demand and create new market opportunities. From a fast neutron-mutagenized population of the waxy corn inbred line HB522, we identified a mutant, *wx-sweet*, whose kernels exhibit both waxy and sweet properties at the filling stage. Using bulked segregant analysis (BSA) and fine mapping, we localized the causal locus to SHRUNKEN1 (*Sh1*) on chromosome 9, which was confirmed by an allelic test with a known Mu-insertion mutant. PCR and sequencing indicated a putative large structural variation in the second and third exons of *Sh1* in *wx-sweet*, likely disrupting the reading frame. This novel sh1 allele significantly reduced sucrose synthase activity. The *sh1* and *waxy1* mutant genes act synergistically to remodel carbohydrate metabolism in *wx-sweet* endosperms. This remodeling, revealed by integrated transcriptomic and metabolomic analyses, drives transcriptional reprogramming and restructures metabolic flux.

These changes thereby enhance sucrose and raffinose accumulation, which underlies the unique waxy-sweet texture in fresh-eating maize endosperm. In summary, our work not provides valuable genetic resources for fresh-eating corn breeding, but also elucidates, for the first time, the molecular mechanism underlying the synergistic formation of waxy and sweet texture.

**Key message:** The novel allele of *Sh1* acts synergistically with *waxy1* to regulate carbohydrate metabolism, which elucidates the molecular mechanism for the unique waxy-sweet texture in fresh-eating corn endosperm. Our work provides new strategies for sweet-waxy maize breeding.

## 1 Introduction

As one of the major crops worldwide, maize provides essential energy via starch accumulated in its kernels. The starch biosynthesis is a complex process orchestrated by a suite of enzymes, such as ADP-glucose pyrophosphorylase (AGP), granule-bound starch synthase (GBSS), soluble starch synthases (SSs), starch branching enzymes (SBEs), starch debranching enzymes (DBEs), and starch phosphorylase (PHO) (Huang et al., 2021; Ajayo et al., 2022). Mutations in the genes encoding these enzymes substantially alter the carbohydrate composition and deposition in the endosperm throughout the grain-filling and milky stages, which in turn determines grain yield and critical quality traits for diverse applications (Botticella et al., 2018).

Waxy corn (*Zea mays* L. sinensis Kulesh) is an important branch variety of maize. It originated from a mutation in waxy gene (*waxy 1*) (Nelson & Rines, 1962). This mutation results in an endosperm starch composition made predominantly of amylopectin. The feature distinguishes it from conventional maize (Nelson & Rines, 1962; Yang, 2008). When cooked,, waxy corns develops sticky, glutinous, and gel-like texture. These qualities are highly valued for its soft and cohesive mouthfeel in both fresh and processed forms (Yang, 2008). In Asia,it is often consumed as fresh ears about 22 days post-pollination (Mehta et al., 2020). Eating quality crucially determines market value.

Sweetness, reflected by soluble sugar content, is a key attribute influencing its sensory appeal. Sweet corn is another specialized type of maize. It is controlled by one or more recessive endosperm mutant genes. Key genes influencing sweetness in maize include *su1*, *sh2*, *bt1*, *bt2*, and *du1* (Chen et al., 2022; Ruanjaichon et al., 2021). The *su1* gene encodes isoamylase, which is responsible for hydrolyzing α-(1→6) glycosidic bonds at aberrant positions in branched polysaccharides (James et al., 1995). Mutation in su1 causes a significant increase in reducing sugars and sucrose content in the kernels. It also leads to substantial phytoglycogen accumulation, which gives sweet corn its desirable taste. As a result, homozygous lines for this gene are widely cultivated as sweet corn (Ledenčan et al., 2022). *Se1* (sugary enhancer1) is a recessive modifier of *su1*. When both genes are double recessive homozygous, (*su1su1se1se1*), the soluble sugar content in kernels can reach approximately twice that of su1 homozygous kernels, holding significant commercial value in modern sweet corn breeding (Zhang et al., 2019). *Sh2* and *bt2* encode the large and small subunits of ADP-glucose pyrophosphorylase (AGPase), respectively. Loss-of-function mutations in these genes significantly reduce AGPase activity and impair ADP-glucose supply (Giroux and Hannah 1994). This causes reduced starch synthesis and substantial increase in soluble sugars, which leads to the super-sweet phenotype (Revilla et al., 2021). Kernels of this type typically exhibit sugar contents of 12% - 22% and an amylopectin proportion below 5%, giving them a crisp and sweet flavor (Pairochteerakul et al., 2018;Chhabra et al., 2022). *Bt1* encodes an adenylate translocator, responsible for transporting cytosolically synthesized ADP-glucose into the amyloplast (Sullivan et al., 1991). Mutations in *bt1*, like in *sh2*, block starch synthesis at a similar stage. Thus, during the milky, *bt1* kernels have higher reducing sugar and sucrose content than *su1* types (Bhave et al., 1990; James et al., 1995). *Du1* encodes soluble starch synthase SSSII. Mutation in this gene also results in significantly higher sugar concentrations during kernel development compared to the wild type (Gao et al., 1998). Based on how mutations affect sugar accumulation, sweet corn production can be categorized into three main types: standard sweet corn (*su1*), enhanced sweet corn (*su1se1*), and super-sweet corn (e.g., types based on *sh2*, *bt2*) (Mehta et al., 2017).

Market demand for fresh-eating corn, such as waxy and sweet corn, is increasing. In China, the annual cultivation area now exceeds 1.67 million hectares, with consumption reaching 75 billion ears (Shi et al., 2019). However, traditional sweet corn and waxy corn varieties cannot simultaneously satisfy consumer preferences for both high sweetness and a strong waxy texture. Studies have shown that crossing commercially available sweet-tasting genes with the waxy gene (*waxy1*) does not produce kernels exhibiting both sweet and waxy sensory characteristics (Ruanjaichon et al., 2022). To overcome this challenge, breeders have developed a novel corn model in which “sweet and waxy kernels coexisting on the same ear”. This approach utilizes the double recessive heterozygous state of sweet and waxy genes and their segregation in progeny, allowing sweet and waxy kernels to appear on a single ear in defined ratios (e.g., 1:3 or 7:9) (Sheng et al., 2023). For instance, by crossing waxy corn with sweet corn, selecting for the waxy-super-sweet genotype (*waxy1waxy1sh2sh2*) in the F2 generation, and subsequently backcrossing with standard waxy corn, results in a novel sweet-waxy corn type. The resulting ears bear both waxy kernels and waxy-sweet kernels in a 1:3 ratio (Wu, 2003).

Although existing commercial corn hybrids featuring a mix of sweet and waxy kernels on the same ear offer flavor variety, The different types exhibited asynchronous dehydration rates and quality deterioration. which narrows the optimal harvest window to just 3–5 days and shortens shelf life, restricting large-scale commercialization (Prai-Anun et al., 2024 ; Fuengtee et al., 2020). To develop novel germplasm featuring both waxy texture and high sweetness within a single kernel, this study utilized the paternal inbred line HB522, derived from China’s registered waxy corn cultivar ‘Su Yunuo 1’, as the starting material. Through fast neutron mutagenesis and screening, we identified a novel genetic material, *wx-sweet*, of which kernels uniquely possess both high sweetness (soluble sugar content is more than 15%) and high waxiness (amylopectin content is more than 95%).

Map-based cloning confirmed that this phenotype is caused by a loss-of-function mutation in the *Sh1* gene. This mutation partially impairs sucrose synthase activity, diverting a portion of carbon substrates towards soluble sugar accumulation. Moreover, compensatory mechanisms involving genes like *Mn1*, *sh2*, and *bt2* allow for the partial preservation of amylopectin biosynthesis, thereby enabling the dual sweet-waxy characteristic (Zhang et al., 2020;Deng et al., 2020). We subsequently developed a functionally linked molecular marker for this allele and applied it in backcross breeding to create a new fresh corn variety with uniform sweet-waxy kernels. The germplasm and genetic insights presented here hold substantial promise for both scientific research and commercial development.

## 2 Materials and Methods

### 2.1 Plant Materials

To develop novel waxy corn germplasm with high sweetness, dry seeds of the elite waxy maize inbred line Hengbai 522 (HB522), which is the paternal parent of the nationally registered cultivar ‘Su Yunuo 1’, were subjected to mutagenesis via fast neutron irradiation. The irradiation was performed at the China Institute of Atomic Energy using a dose of 3.99 Gy. The 30,000 treated M_0_ seeds were subsequently sown at the Luhe Experimental Station of the Jiangsu Academy of Agricultural Sciences to generate the self-pollinated M1 population. Through three generations of subsequent self-pollination and trait screening, a genetically stable mutant with concurrent sweet and waxy phenotypes in the kernels was identified and named *wx-sweet*.

### 2.2 Phenotypic Analysis of *wx-sweet*

Phenotypic comparison between the wildtype (*wx*) and mutant (*wx-sweet*) was performed using kernels harvested from mature segregating ears. The kernels were first assessed for visual traits, size, internal structure, and the developmental status of the embryo and endosperm. Seed germination capacity was also evaluated. For quantitative measurements, a hundred-kernel weight assay was performed. Three biological replicates were used, each comprising 100 randomly sampled kernels from a distinct segregating ear. Following this, the weights of embryo and endosperm were determined by manually dissecting ten seeds from each genotype. This procedure was repeated three times to generate mean values for embryo and endosperm weights from each replicate.

### 2.3 Endosperm Physiological Traits of *wx-sweet*

Endosperms were isolated from *wx* and *wx-sweet* kernels harvested at various developmental stages (14, 17, 20, 23, 26 DAP) and at maturity. Samples were dried at 75°C until constant weight achieved, ground into a fine powder using a tissue lyser, and sieved (100-mesh). This powder served for all downstream physiological analyses, each conducted with a minimum of three replicates.

**Total Starch Content**: We quantified total starch using the Solarbio Kit (BC0705). Briefly, soluble sugars were extracted with 80% ethanol. The residual starch was acid-hydrolyzed to glucose, which was then measured via the anthrone colorimetric method to calculate starch content (Viles and Silverman, 1949; Clegg, 1956).

**Amylopectin Content**: Using the Solarbio Kit (BC4270), we measured amylopectin content colorimetrically based on its specific reddish-purple complex formation with iodine, following soluble sugar removal by ethanol.

**Soluble Sugar Content**: Total soluble sugars were directly determined by the anthrone method using the Solarbio Kit (BC0035), adhering to the provided protocol (BUYSSE and MERCKX 1993).

**Sucrose Content**: Sucrose was assayed with the Solarbio Kit (BC2465). After alkaline removal of reducing sugars, sucrose was hydrolyzed. The released fructose was reacted with resorcinol, and the absorbance at 480 nm was measured to calculate sucrose concentration (Fils-Lycaon et al., 2011).

### 2.4 Scanning Electron Microscopy

Mature *wx* and *wx-sweet* kernels from the same ear were dried at 42°C for 72 hours. Samples were then longitudinally sectioned with a sharp blade to expose a natural internal surface. After gold coating via an E-100 ion sputter, the samples were imaged using a ZEISS EVO-LS10 scanning electron microscope to analyze surface and cross-sectional ultrastructure.

### 2.5 Assay of Sucrose Synthase Activity in Developing Maize Endosperm

Endosperms from *wx* and *wx-sweet* kernels at different developmental stages (14, 17, 20, 23, and 26 DAP) were separately collected, immediately frozen in liquid nitrogen, and stored at -80°C until analysis. For the assay, a minimum of 20 endosperms were ground into a fine powder in a steel jar pre-cooled with liquid nitrogen using a tissue lyser. Approximately 50 mg of the resulting powder was accurately weighed into a pre-cooled centrifuge tube for sucrose synthase (SUS) activity determination. SUS activity was measured using the Solarbio Sucrose Synthase Assay Kit (BC0580) according to the manufacturer’s protocol, based on a visible spectrophotometric method. The enzyme catalyzes the reaction of fructose and UDP-glucose to produce UDP and sucrose(Schrader and Sauter, 2002). The synthesized sucrose then reacts with resorcinol to generate a colored product with a characteristic absorption peak at 480 nm, the absorbance of which is proportional to the enzyme activity. Each sample was assayed with three technical replicates, and the absorbance at 480 nm was recorded using a Beckman Coulter DU730 UV-Vis spectrophotometer.

### 2.6 Sugar Analysis

Endosperms from *wx* and *wx-sweet* kernels at various developmental stages (14, 17, 20, 23, and 26 DAP) were collected for targeted sugar metabolite profiling, which was conducted by MetWare Co., Ltd. Sugar extraction was performed following the method described by Chen (2022). The extracts were analyzed using an Agilent 7890B-5977B GC-MS system equipped with a DB-5MS capillary column (30 m × 0.25 mm × 0.25 μm, J&W Scientific). For each compound, multiple characteristic peak areas were weighted and merged into a single abundance value. All data were mean-centered and auto-scaled prior to one-way analysis of variance in R. Resulting *P*-values were adjusted using the Bonferroni–Hochberg method.

Maize endosperm from *wx* and *wx-sweet* kernels were freeze-dried and ground into powder using a grinding mill (30 Hz, 1.5 min). Exactly 20 mg of the powder was weighed and mixed with 500 μL of a methanol:isopropanol:water (3:3:2, V/V/V) extraction solution. The mixture was vortexed for 3 min and then subjected to ultrasonic treatment in an ice-water bath for 30 min. Subsequently, the mixture was centrifuged at 4°C and 14,000 r/min for 3 min. A 50 μL aliquot of the supernatant was collected, mixed with 20 μL of an internal standard solution (concentration: 1000 μg/mL), and then dried using nitrogen blow-down followed by freeze-drying. Afterward, 100 μL of methoxyamine hydrochloride in pyridine (15 mg/mL) was added to the freeze-dried sample, and the mixture was incubated at 37°C for 2 h. Then, 100 μL of BSTFA was added, and the derivatization was continued at the same temperature for 30 min to obtain the derivatized solution. Finally, 50 μL of the derivatized solution was diluted to 1 mL with n-hexane, transferred into a brown injection vial, and prepared for GC-MS analysis. The GC-MS analysis was performed on an Agilent 8890 gas chromatography system coupled with a 5977B mass spectrometer, using a DB-5MS capillary column (30 m × 0.25 mm × 0.25 μm, J&W Scientific, USA). The analytical method referred to the study by Luo (2021) and Chen (2022). For each metabolite, the corresponding multiple diagnostic ions were consolidated into a single abundance value through weighted integration. Prior to performing one-way ANOVA in R, all datasets were subjected to mean centering and autoscaling normalization. The resulting *P*-values were subsequently corrected via the Bonferroni-Hochberg procedure.

### 2.7 Pasting Property Analysis (RVA)

Endosperm at different developmental stages (14, 17, 20, 23, 26 DAP) and maturity from *wx* and *wx-sweet* kernels were collected, freeze-dried using a vacuum freeze-dryer (FD-1, Biocool), ground into powder, and sieved. Precisely 5.0 g of the resulting powder was mixed with 25.0 g of distilled water in a dedicated RVA aluminum canister. The pasting properties were then determined using a Rapid Visco Analyzer (TecMaster, Perten) following Standard Method 1. Each sample was analyzed in triplicate, and the data were processed using the accompanying TCW software (Castanha et al., 2021).

### 2.8 BSA-seq Analysis

An F₂ segregation population was generated by self-pollinating F₁ plants derived from a cross between the waxy maize inbred line HB522 and the *wx-sweet* mutant. Genomic DNA was extracted from 20 *wx* and 20 *wx-sweet* seedlings using the CTAB method and subjected to genome resequencing. Using the B73 (V4) reference genome (Jiao et al., 2017), SNP calling was performed with GATK v4.1.1.0. Initial filtering was applied with the following thresholds: QD < 2.0, QUAL < 30.0, MQ < 40.0, FS > 60.0, SOR > 3.0, MQRankSum < −12.5, and ReadPosRankSum < −8.0. Subsequent filtering using PLINK v1.90 retained SNPs with a missing rate < 0.2 and a minor allele frequency > 0.05.

SNP-index calculation and QTL mapping were then conducted using the QTLseqr package( ). The filtered SNP dataset was imported via the importFromTable function, with wx samples designated as the highBulk group and *wx-sweet* samples as the lowBulk group. The runQTLseqAnalysis (Mansfeld et al., 2018) function was applied with the population type set as F₂ and bulk sizes specified as c (20, 20) to quantify allele frequency differences between the two groups. Results were visualized using the R software, and genomic regions with |ΔSNP-index| > 0.4 were identified as candidate intervals.

### 2.9 Fine Mapping and Allelic Complementation Test

Based on the high-differentiation region identified in the BAS-seq analysis, Indel markers (Supplementary Table S5) were developed within the target interval using the HB522 × *wx-sweet* F₂ population. Polymorphic markers between the parents were selected for genotyping *wx-sweet* individuals. Recombinant events were statistically analyzed to refine the candidate gene region. Genes within the mapped interval and their functional annotations were retrieved from the MaizeGDB database.

To determine whether *wx-sweet* is an allele of *sh1*, a complementation test was performed by crossing *wx-sweet* with the *sh1*-Mu mutant. An F₁ kernel exhibiting the mutant phenotype would indicate allelism, while a wild phenotype would suggest non-allelism.

### 2.10 Candidate Gene Sequencing

Gene-specific primers (Supplementary Table S5) were designed based on the sequence of the candidate gene *sh1* from the B73 reference genome. PCR amplification was performed using high-fidelity DNA polymerase with genomic DNA from *wx* and *wx-sweet* as templates, and the products were sequenced. Simultaneously, full-length *sh1* cDNA was amplified and sequenced using cDNA templates derived from kernels at 14 DAP. The PCR reaction mixture consisted of 25 μL of 2× Super Pfx Master Mix, 1.5 μL of genomic DNA, 2.5 μL each of forward and reverse primers, and 18.5 μL of ddH₂O. The PCR program was as follows: initial denaturation at 98°C for 30 s; 35 cycles of 95°C for 10 s, 58°C for 10 s, and 72°C for 20 s; and a final extension at 72°C for 5 min. Amplified products were separated by 2% agarose gel electrophoresis, followed by sequencing and sequence alignment analysis.

### 2.11 Development and Genotyping of a Functional Marker for *wx-sweet*

A *wx-sweet*-specific molecular marker was developed by designing primers (Supplementary Table S5) targeting the sequence of the second intron of the *sh1* gene using Primer 5.0. PCR amplification was conducted using DNA from *wx* and *wx-sweet* as templates at an annealing temperature of 58°C and an extension time of 40s. The amplification products were resolved and detected via 3% agarose gel electrophoresis.

### 2.12 RNA Extraction and qRT-PCR Analysis

Total RNA was extracted from *wx* and *wx-sweet* kernels at different developmental stages (14, 17, 20, 23, 26 DAP) using the SV Total RNA Isolation System. One microgram of total RNA was treated with gDNA Eraser and reverse-transcribed using the PrimeScript™ RT reagent Kit. qRT-PCR was performed on a Roche Light Cycler 2.0 system using SYBR Premix Ex Taq™, following the method of Ning (2023). The 10 μL reaction mixture contained 5 μL of SYBR Premix, 0.8 μL each of 10 μM forward and reverse primers, and 1 μL of cDNA, made up to volume with nuclease-free water. Primer sequences are listed in Supplementary Table S5. The Ubiquitin gene was used as an internal control, and the relative expression levels of target genes were calculated using 2^−ΔΔCt^ method (Livak and Schmittgen, 2001). Three biological replicates were included for each sample.

### 2.13 RNA-seq data analysis

Endosperm of *wx* and *wx-sweet* maize kernels were collected at five developmental stages (14, 17, 20, 23, and 26 days after pollination, DAP), with three biological replicates per time point. Total RNA from these samples was sent to Berry Genomics Co., Ltd. for RNA sequencing. The libraries were constructed and sequenced on an Illumina HiSeq 2500 platform to generate 150 bp paired-end reads.

Raw read quality was assessed with FastQC (De Sena Brandine and Smith, 2019). Subsequently, reads were trimmed to 100 nucleotides using the fastx_trimmer tool from the FASTX Toolkit (http://hannonlab.cshl.edu/fastx_tool kit/index.html) to remove low-quality bases. Only reads longer than 50 bp after trimming were retained for subsequent analysis.

High-quality reads were aligned to the B73 reference (V4) maize cDNA sequences (downloaded from www.maizegdb.org) and transcript abundances were quantified as TPM (Transcripts Per Million) using Salmon (v1.1.0) (Schurch et al., 2016). Differential expression analysis was performed using DESeq2, identifying genes with an absolute |log_2_ FoldChange| ≥ 1 and *P*-value < 0.05 as statistically significant differentially expressed genes (DEGs). Finally, Gene Ontology (GO) enrichment analysis for the identified DEGs was conducted using the GOseq R package.

### 2.14 Statistical Analysis

Statistical analysis was performed using GraphPad Prism 5. An unpaired two-tailed Student’s *t*-test was applied. Data are presented as mean ± standard error (SE). Significance levels were set at *P* < 0.05, and *P* < 0.01.

## 3 Results

### 3.1 Screening for *wx-sweet* by fast-neutron mutagenesis

Through fast neutron mutagenesis of seeds from the elite waxy maize inbred line HB522, followed by three consecutive generations of self-pollination and phenotypic screening, a unique mutant ear, designated *wx-sweet*, was identified among 1,203 M₃ ears. The progeny kernels of this ear exhibited phenotypic segregation, with the ratio of normal, fully filled kernels to shrunken kernels fitting a 3:1 ratio (Fig. 1A, B). Longitudinal sectioning of the mutant kernels revealed defects in endosperm development (Fig. 1C). 100-kernel weight analysis indicated a 18.72% reduction in mutant (*wx-sweet*) kernels compared to the wild type (*wx*) (Fig. 1F). Further measurement showed that the endosperm accounted for only 64.95% of the mutant kernel weight (Fig. 1H). Germination assays demonstrated no significant difference in germination rate between the mutant and wild type (Fig. 1D, G). The shrunken kernels germinated normally, and subsequent seedling growth showed no observable abnormalities (Fig. 1E).

**Figure 1.**
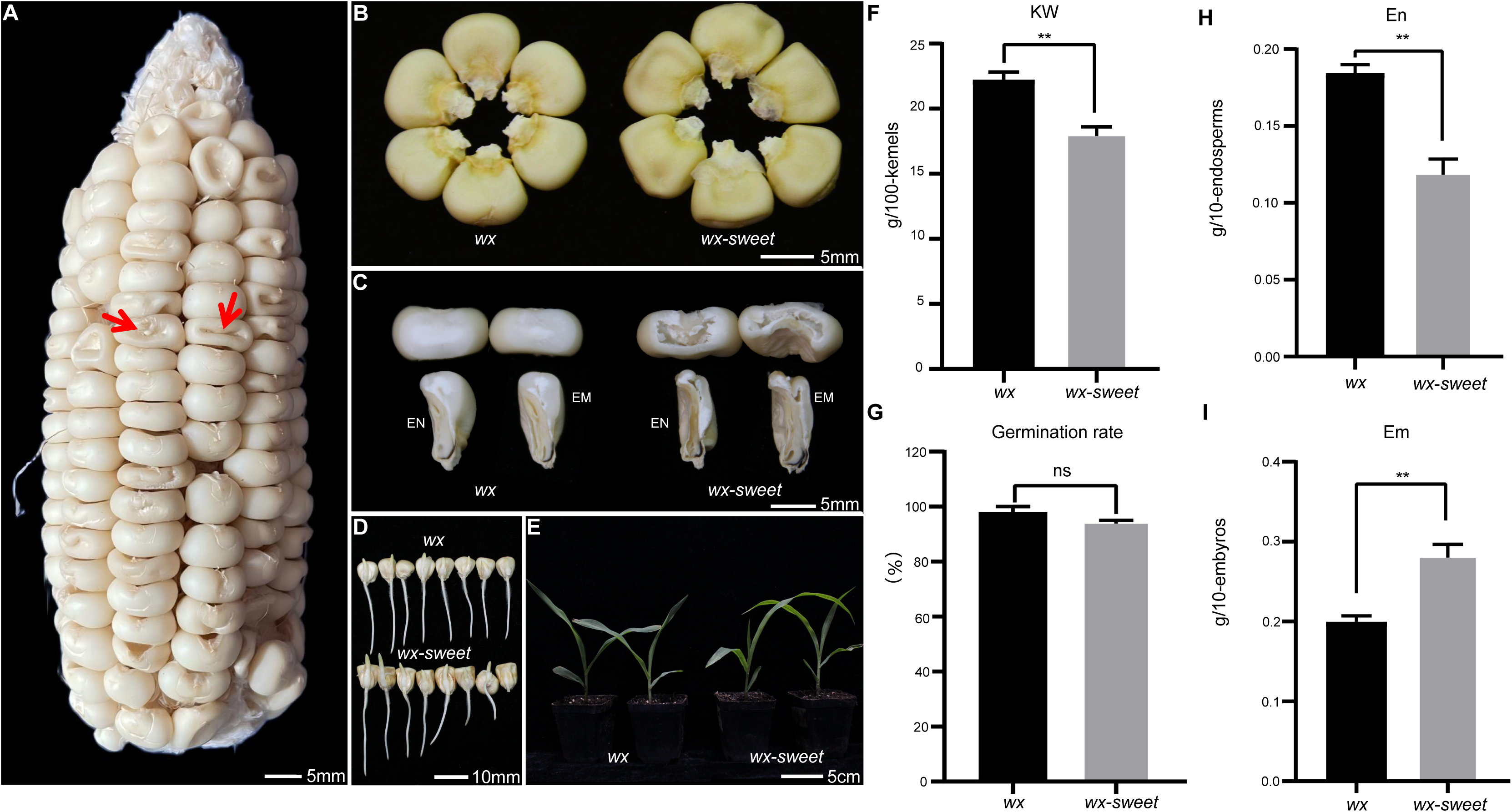
Phenotypic Features of *wx* and *wx-sweet* in the HB522 Background. A: Mature ear of the F2 (*wx* × *wx-sweet*) segregating population. Red arrowheads indicate typical *wx-sweet* kernels. B: Wild-type (*wx*) and *wx-sweet* mature kernels viewed under natural light. C: Transverse and longitudinal sections of kernels; EM, embryo; En, endosperm. Scale bar = 5mm. D: Germination status of *wx* and *wx-sweet* kernel, scale bar = 1cm. E: Comparison of seedlings between *wx* and *wx-sweet*, scale bar = 5cm. F: 100-kernel weight of *wx* and *wx-sweet* mutant kernels in the F_2_ segregating population; G: Germination rate of *wx* and *wx-sweet*. H-I: Comparison of embryos (H) and endosperms (I) between *wx* and *wx-sweet*. Note: Data are presented as mean ± SD from independent replicates; F–I were repeated three times. Bar graphs represent mean ± standard error (SE). *: Significant difference at *P* < 0.05; **: Significant difference at *P* < 0.01. g: gram. Em: Embryo; En: Endosperm.

Analysis of kernel composition revealed significant reductions in both total starch and amylopectin contents in the mutant (*wx-sweet*), which were 77.85% and 82.26% of wild type (*wx*) levels, respectively (Fig. 2A, C; Supplementary Fig. S1A). The proportion of amylopectin relative to total starch remained high at 95.6%. Notably, the soluble sugar content in the mutant kernels increased markedly to 15.03 (mg/g), representing a 2.82-fold increase over the *wx* (Fig. 2B). Meanwhile, sucrose content in the *wx-sweet* kernels also increased significantly (Fig. 2D). Scanning electron microscopy of mature kernels showed that, compared to the wild type, the *wx-sweet* mutant exhibited smaller starch granules with irregular arrangements and the presence of small surface pits (Fig. 2E–H).

**Figure 2.**
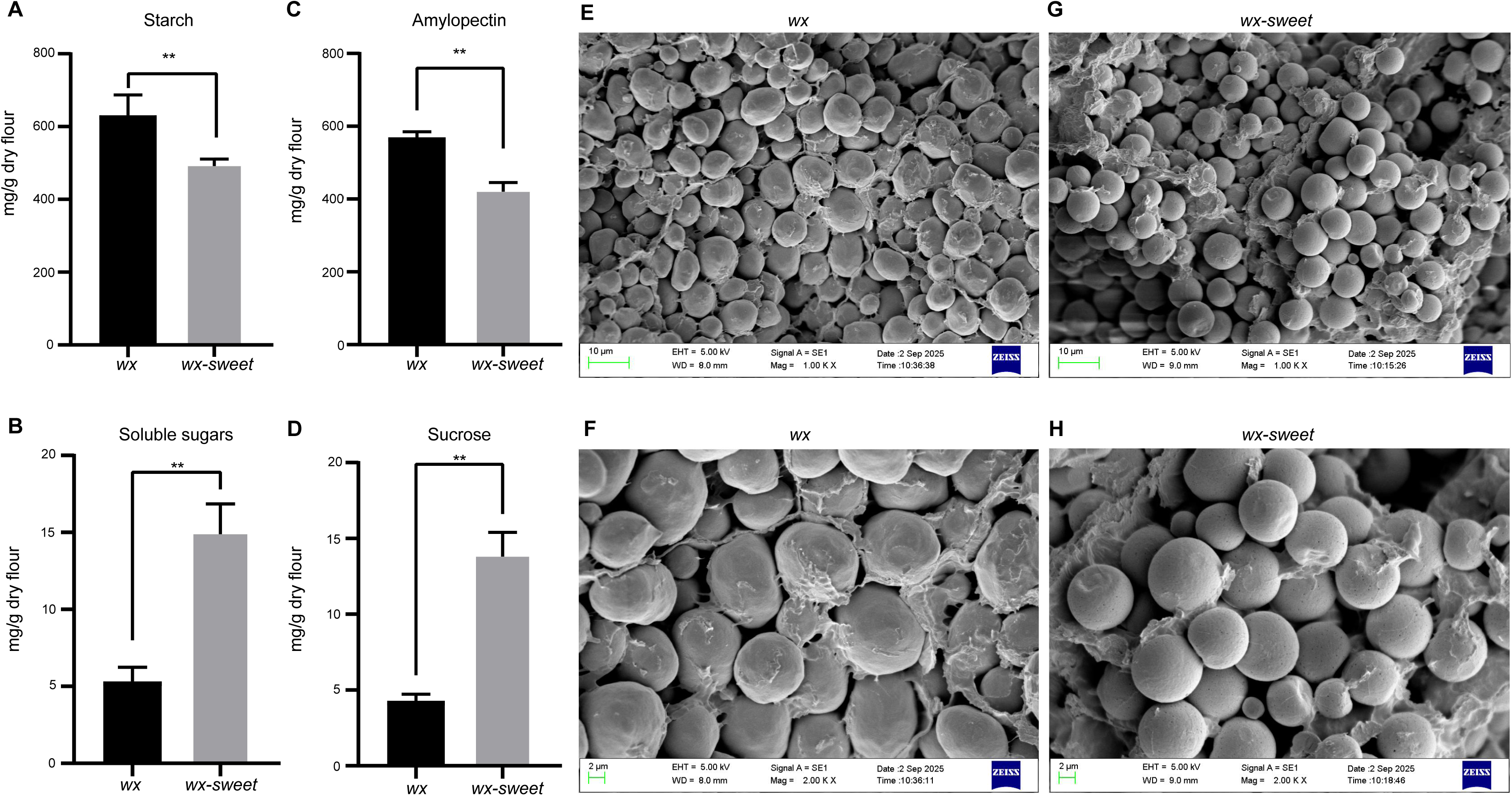
Biochemical Analysis and Electron Microscopy Observation of *wx* and *wx-sweet* Mutant. A: Starch content in mature kernels of *wx* and *wx-sweet* mutants. B: Soluble sugar content in mature kernels of *wx* and *wx-sweet* mutants. C: Amylopectin content in mature kernels of *wx* and *wx-sweet* mutants. D: Sucrose content in mature kernels of *wx* and *wx-sweet* mutants. E-F: Scanning electron microscopy of the central regions of the mature endosperm of *wx* (E) and *wx-sweet* (F) kernel. SEM ×100K, bar = 10 µm; G-H: Scanning electron microscopy of the central regions of the mature endosperm of *wx* (G) and *wx-sweet* (H) kernel. SEM ×200K, bar = 2 µm; Note: Experiments were repeated three times. Bar graphs represent mean ± SE. *: Significant difference at *P* < 0.05; **: Significant difference at *P* < 0.01.

Collectively, these results indicate that the loss of *wx-sweet* gene function disrupts the normal processes of sugar metabolism and starch accumulation in the maize kernel endosperm, leading to impaired starch synthesis, substantial soluble sugar accumulation.

### 3.2 Cloning of *wx-sweet*

Genetic analysis revealed that the F₁ progeny from a cross between *wx-sweet* and HB522 exhibited a wild type phenotype. Self-pollination of the F₁ plants yielded an F₂ population that segregated for kernel phenotypes. Among 170 F₂ kernels, 123 displayed the wild phenotype and 47 were mutant, fitting a 3:1 ratio (*χ²* = 1.27, *P* > 0.05), indicating that the *wx-sweet* mutant phenotype is controlled by a single recessive nuclear gene.

To map the candidate gene, we selected 20 wildtype and 20 mutant plants from the F₂ population for whole-genome resequencing. We performed a genome-wide analysis of SNP-index changes for all polymorphic SNP loci (Mansfeld et al., 2018), which revealed an interval on chromosome 9 (4.91-19.83 Mb) that may be linked to the target trait (Fig. 3A). Within this region, InDel markers were developed. Genotyping of 170 recessive F₂ individuals initially delimited the candidate region between markers ID2 and ID3 (10.19-13.62 Mb). By expanding the F₂ population to 1,241 individuals and conducting further genotyping, the candidate interval was finely mapped to a 634-kb region (10.83-11.47 Mb), flanked by ID22 and ID24 (Fig. 3B).

**Figure 3.**
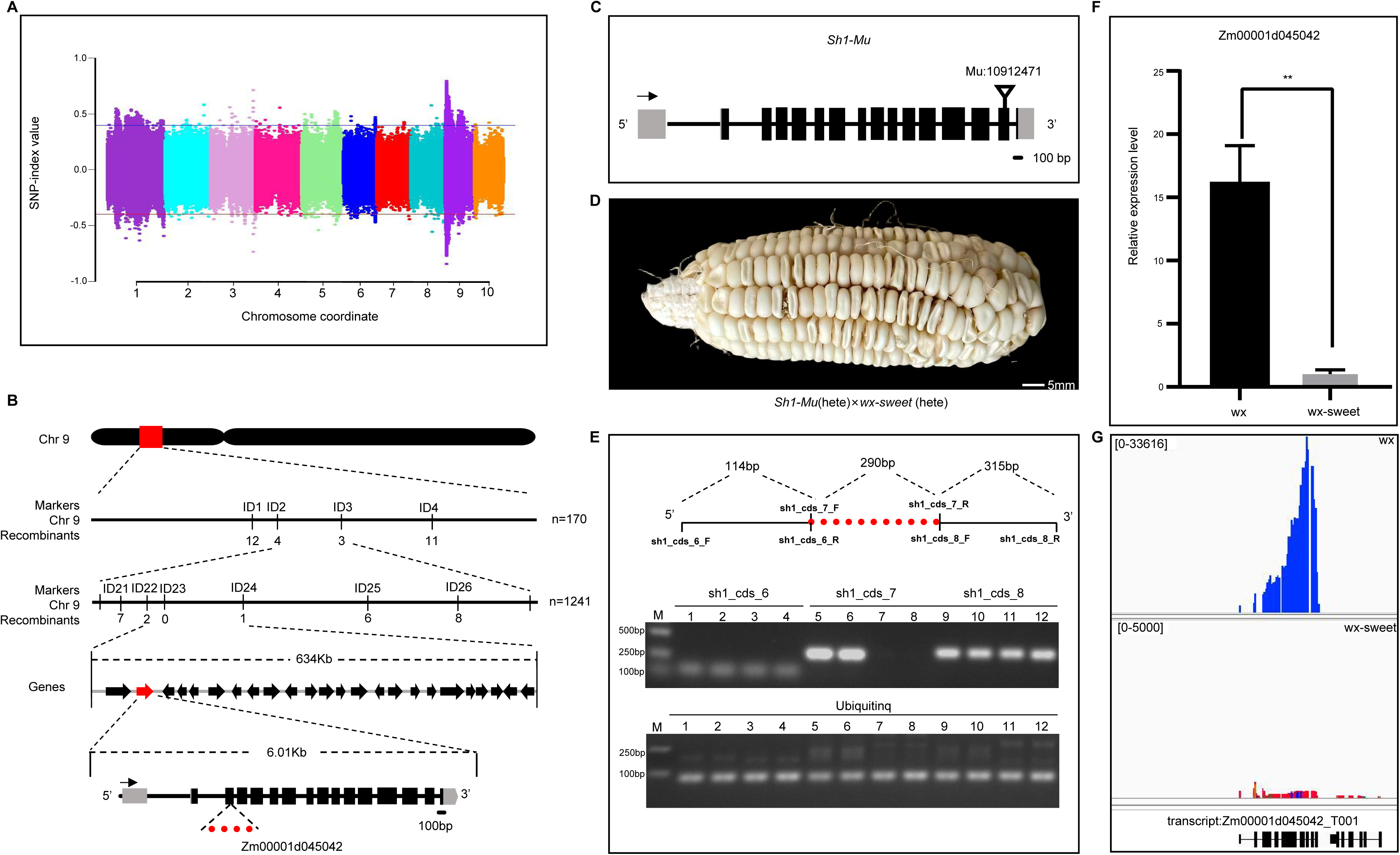
Map-Based Cloning and Identification of *wx-sweet*. A: BSA-seq analysis mapped the *wx-sweet* gene to chromosome 9. B: Linkage markers ID1–ID4 identified by genotyping 170 *wx-sweet* individuals; Fine mapping of wx-sweet using 1,241 individuals from the F_2_ population. *wx-sweet* was localized to a 634-kb region between markers ID22 and ID24. Marker names are indicated above the black vertical solid lines, and the number of recombinant individuals identified by each marker is shown below; N: Number of individuals used for mapping; Recombinant: Number of recombinant individuals; C: Gene structure of *Sh1* and insertion site of the *Sh1*-Mu mutant; D: An allelic complementation test between *sh1*-Mu (heterozygous) and *wx-sweet* (heterozygous). Hybrid ears (*sh1*-Mu/+ × *wx-sweet*/+) consistently segregated approximately one quarter mutant kernels, scale bar = 5 mm; E: PCR amplification of the *Sh1* gene using cDNA from *wx* and *wx-sweet* kernels. See Supplementary Figure S2 for additional details. F: qRT-PCR analysis of *Sh1* expression in *wx* and *wx-sweet* kernels at 14 DAP ; G: Comparison of *Sh1* reads in RNA-seq data from 14 DAP endosperms of *wx* and *wx-sweet* kernels.

According to MaizeGDB annotations, this interval contains 24 open reading frames (Supplementary Table S1), 14 of which are expressed in seeds. Functional annotation of these candidates identified Zm00001d045042, which encodes sucrose synthase *Sh1*, a key enzyme catalyzing the cleavage of sucrose into fructose and UDP-glucose during grain filling. Loss-of-function mutations in *Sh1* are known to cause a shrunken kernel phenotype (Chourey and Schwartz), consistent with the *wx-sweet* mutant. Thus, we conducted an allelic complementation test to determine whether *wx-sweet* is allelic to *Sh1*. We identified a mutant from the ChinaMu population carrying a MuA2 insertion +5,538 bp of the *sh1* (Liang et al., 2019). After backcrossing this mutant into the HB522 background (Zhou et al., 2025), we obtained an *sh1*-Mu allele in the HB522 genetic background (Fig. 3C). *wx-sweet* (heterozygote) were then crossed with *sh1*-Mu (heterozygote), and the F_1_ ears consistently segregated approximately one-quarter mutant kernels (Fig. 3D). These results confirm that *wx-sweet* is an allelic mutant of the *Sh1* gene.

### 3.3 Identification of the *Sh1* Mutation and Analysis of Its Expression

To identify potential variations in the *Sh1* gene in the *wx-sweet* mutant, we designed primers based on the *Sh1* sequence obtained from second-generation sequencing data of the *wx* genome and performed amplification and sequencing of the gene in both *wx* and *wx-sweet* kernels (Supplementary Table S5). When PCR amplification was performed using primers spanning the region between the second and third exons, no target fragment was amplified from either genomic DNA or cDNA of endosperm at 14 DAP (Fig. 3E; Supplementary Fig. S2), whereas the expected fragments were successfully amplified from both DNA and cDNA templates of the *wx* (Fig. 3E; Supplementary Fig. S2). This results suggest that a large structural variation may exist between the second and third exons in the *sh1* allele of *wx-sweet*. This variation may lead to an altered reading frame of the *Sh1* gene in. Further Sanger sequencing results revealed 21 nucleotide variations and one two-nucleotide deletion within the *Sh1* open reading frame of compared with *wx*, in addition to the non-amplifiable gap (Fig. 3E; Supplementary Table S2).

To determine the effect of these variations on the transcriptional level of *Sh1*, we measured its transcript abundance in the endosperms of *wx* and *wx-sweet* kernels at 14 DAP by quantitative RT-PCR. The results showed a significant reduction in *Sh1* expression of *wx-sweet* (Fig. 3F). This finding was corroborated by subsequent transcriptome analysis. Visualization of the RNA-seq read alignments across the *Sh1* locus using IGV revealed drastically reduced read coverage in *wx-sweet*, appearing as a faint peak (Fig. 3G). In stark contrast, the *wx* sample exhibited abundant read enrichment and a sharp, high peak spanning the *Sh1* gene region (Fig. 3G). Together, these results strongly indicate that transcription of the *Sh1* gene is severely suppressed in *wx-sweet* kernels.

### 3.4 The *sh1* Mutation Significantly Reduces Sucrose Synthase Activity in Endosperms

This enzymatic pattern was consistent with changes in the expression of SUS-encoding genes. In maize, three genes reported (*Sh1*, *Sus1*, and *Sus2*) encode sucrose synthases, with *Sh1* being the major contributor to SUS activity in the endosperm (Deng et al., 2020). To investigate the impact of the *sh1* mutation on sucrose synthase (SUS) activity in the *wx-sweet* mutant, we measured total SUS activity in endosperms of *wx* and *wx-sweet* at five developmental stages. The results showed a similar trend in both genotypes, characterized by an initial increase followed by a decrease, with peak activity occurring around 17 DAP (Fig. 4A). However, SUS activity was significantly lower in *wx-sweet* than in *wx* across all developmental stages. Notably, the rate of decline in total SUS activity during later development was slower in *wx-sweet* compared to *wx*.

**Figure 4.**
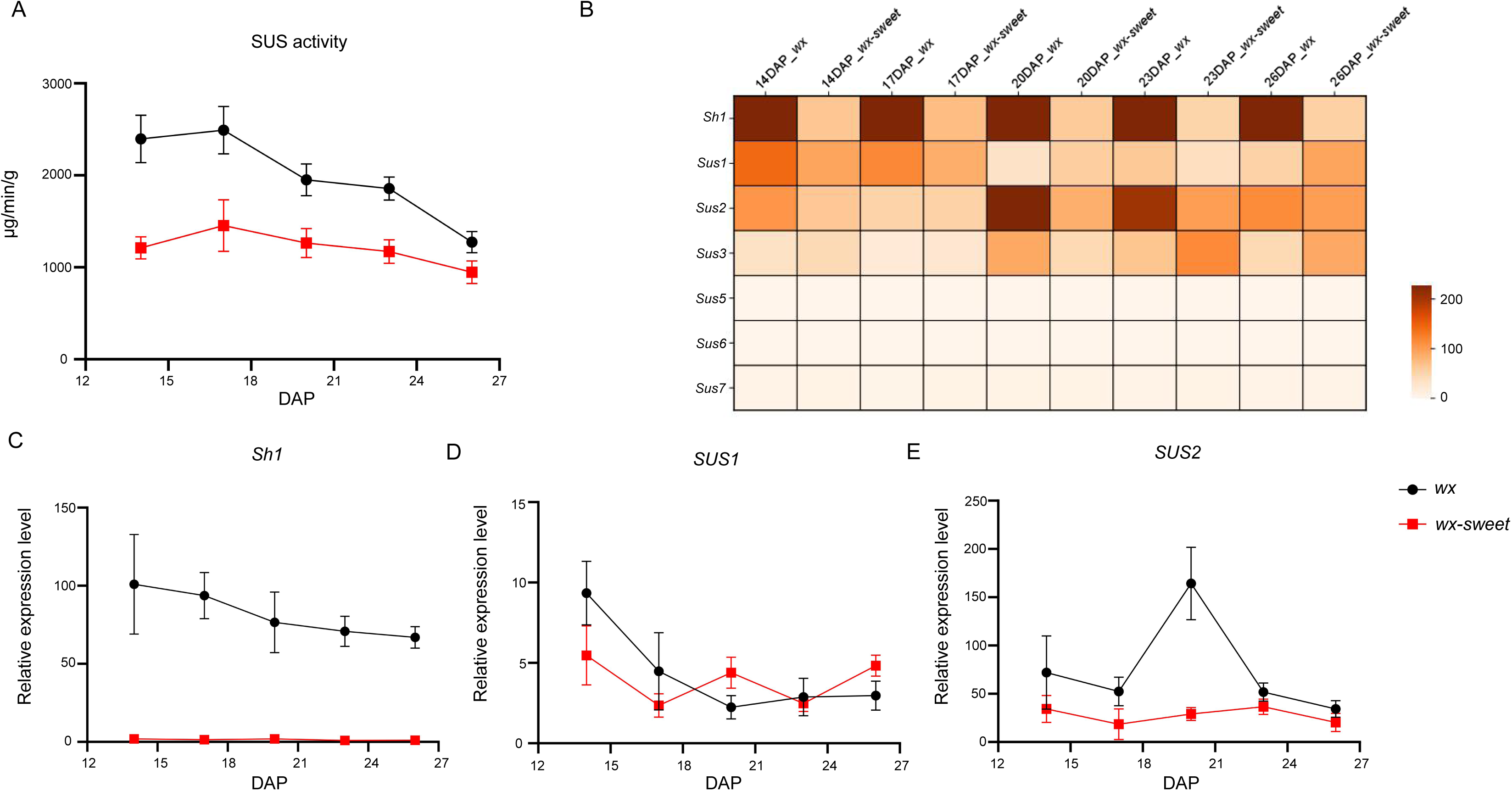
Total SUS Activity and Transcript Levels of Sus Genes in Developing Endosperms of *wx* and *wx-sweet*. A: Total SUS activity in endosperm of *wx* and *wx-sweet* at different development stage (14, 17, 20, 23, and 26 DAP). Data represent mean ± SD from three replicates. One unit of enzyme activity (μg min⁻¹ g⁻¹) is defined as the amount catalyzing the formation of 1 μg sucrose per gram of tissue per minute. B: TPM values of *Sh1*, *SUS1*, and *SUS2* genes in *wx* and endosperm at different development stage (14, 17, 20, 23, and 26 DAP). C-E: Transcript levels of *Sh1*, *SUS1*, and *SUS2* genes in endosperms of *wx* and *wx-sweet* at different development stage (14, 17, 20, 23, and 26 DAP), normalized to Ubiquitin. Data represent mean ± SD from three replicates. * and ** indicate significant differences compared to *wx* at *P* < 0.05 and *P* < 0.01, respectively (Student’s *t*-test).

qRT-PCR analysis of these SUS-encoding genes in *wx* and *wx-sweet* endosperms across the five developmental stages revealed a substantial reduction in *Sh1* transcript levels in *wx-sweet* (Fig. 4B,C). In contrast, *Sus1* expression was significantly upregulated in *wx-sweet* at 20 and 26 DAP compared with *wx* (Fig. 4D), while *Sus2* expression showed a slight overall decrease in the mutant (Fig. 4D). RNA-seq analysis corroborated the qRT-PCR results, confirming these expression patterns. Totally, the mutation in *Sh1* and its consequent transcriptional downregulation are the primary causes of the significant reduction in total SUS activity. The upregulation of other SUS-encoding genes at later developmental stages may partially account for the slower decline in total SUS activity observed in the *wx-sweet* endosperm (Fig. 4B).

### 3.5 Transcriptomic Profiling Reveals Reprogramming of Carbohydrate Metabolism Gene Expression in Endosperms by *sh1*

To detect the transcriptional regulatory impact of the *sh1* mutation on endosperm development, RNA-seq analysis was performed on endosperms of *wx* and *wx-sweet* mutants at five developmental stages (14, 17, 20, 23, and 26 DAP). After quality control, an average of 87.8% of the sequencing reads were uniquely mapped to the B73 (V4) reference transcriptome. Following the removal of low-expression transcripts (total counts < 10), an average of 22,793 genes per stage were retained for subsequent analysis across the five developmental stages (Supplementary Dataset S1). Differential expression analysis between *wx* and *wx-sweet* at each time point was conducted using DESeq2 (criteria: |log₂FoldChange| > 1, *P*-value < 0.05). The number of differentially expressed genes (DEGs) was lowest at 14 DAP, with 1,172 up-regulated and 1,084 down-regulated genes in *wx-sweet* (Fig. 5A; Supplementary Dataset S2), and highest at 23 DAP, with 3,399 up-regulated and 1,657 down-regulated genes. The number of DEGs increased markedly during endosperm development, rising from 2,256 at 14 DAP to 4,470 at 26 DAP. Among these, 403 DEGs were consistently identified across all five stages (Fig. 5B).

**Figure 5.**
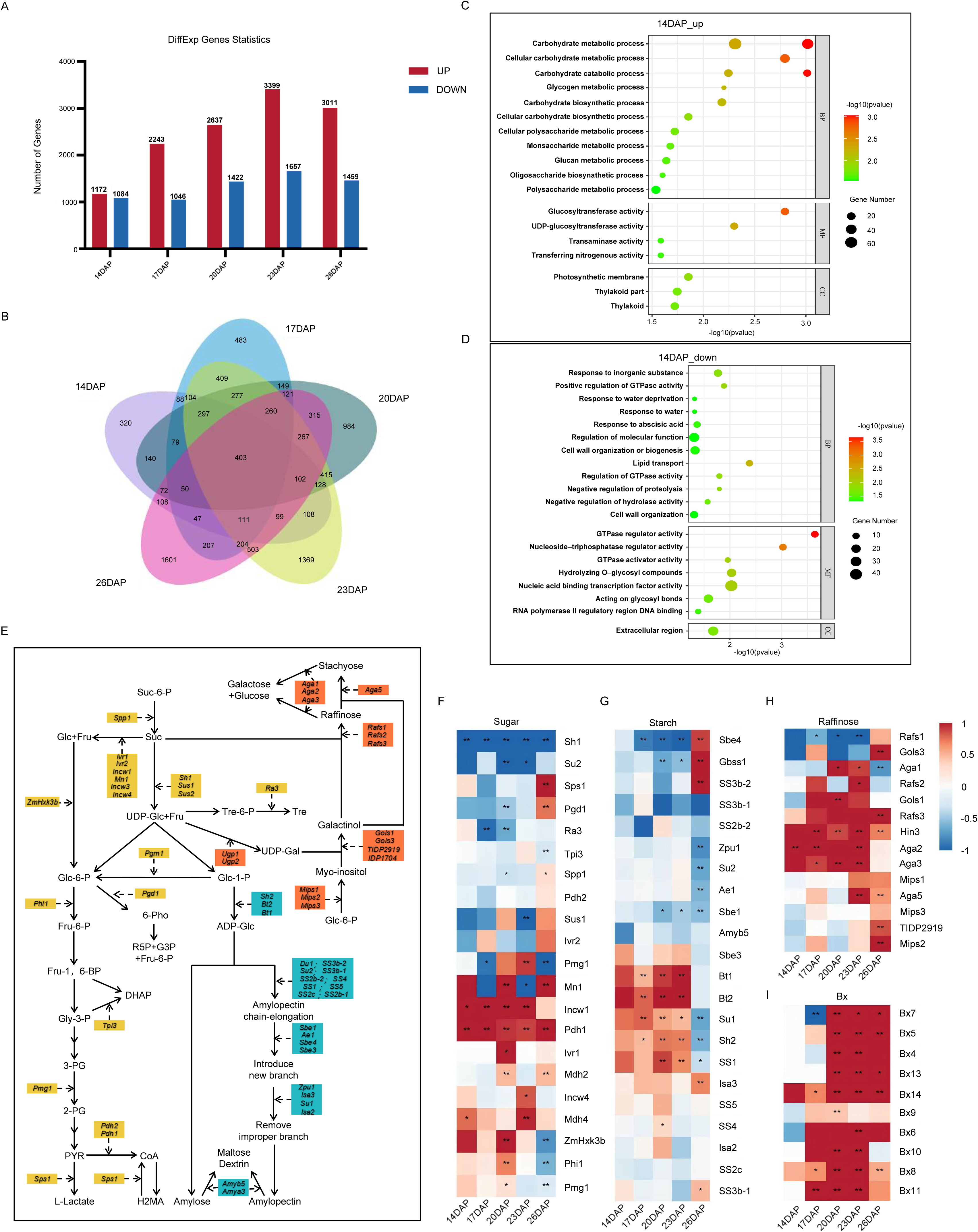
Transcriptome Comparison of *wx* and *wx-sweet* Endosperm. A: The number of differentially expressed genes (DEGs) in *wx* and *wx-sweet* endosperm across different developmental stages (14, 17, 20, 23, and 26 DAP), DEGs were identified using DESeq2 with the criteria of |log2FoldChange| ≥ 1 and *P*-value < 0.05; B: Venn diagram of *sh1*-regulated DEGs at different development stage (14, 17, 20, 23, and 26 DAP); C: GO enrichment of upregulated DEGs in *wx-sweet* vs *wx* kernels at 14 DAP; D: GO enrichment of downregulated DEGs in *wx-sweet* vs *wx* kernels at 14 DAP; E: Schematic of sugar, raffinose, and starch metabolic pathways in maize kernels. Genes involved in sugar metabolism are highlighted in yellow, raffinose-related genes in orange, and starch synthesis-related genes in blue. F: Heatmaps of log₂FoldChanges for genes involved sugar metabolic pathways in maize; G: Heatmaps of log₂FoldChanges for genes involved starch metabolic pathways in maize; H: Heatmaps of log₂FoldChanges for genes involved raffinose metabolic pathways in maize; I: Heatmaps of log₂FoldChanges for maize benzoxazinoid pathway genes detected in this study. The heatmaps display values (*wx-sweet* vs *wx*) from 14, 17, 20, 23, and 26 DAP, which were used for clustering. See Supplementary Dataset S1 and S3 for further details. Note : Raf, Raffinose; 2-PG,2-Phosphoglycerate; 3-PG, 3-Phosphoglycerate; 6-Pho, 6-Phosphogluconolactone; ADP-Glc, ADP-Glucose; CoA, Coenzyme A; DHAP, Dihydroxyacetone phosphate; Fru, Fructose; Fru-1,6-BP, Fructose-1,6-Bisphosphate; Fru-6-P, Fructose-6-Phosphate; Glc,Glucose; Glc-1-P, Glucose-1-Phosphate; Glc-6-P, Glucose-6-Phosphate; G3P, Glyceraldehyde-3-Phosphate; H2MA, malic acid; PYR, Pyruvate; R5P, Ribose-5-Phosphate; Sta,Stachyose; Suc, Sucrose; Suc-6-P, Sucrose-6-Phosphate; Tre, Trehalose; Tre-6-P, Trehalose-6-Phosphate; UDP-Gal, Uridine diphosphate galactose; UDP-Glc, Uridine diphosphate glucose; SS3b-2, starch synthase3; Sbe4, starch branching enzyme4; Zpu1, pullulanase-type starch debranching enzyme1; Amya3, alpha amylase3; Sh2, shrunken2; Su1, sugary1; Bt2, brittle endosperm2; SS5, starch synthase5; SS2b-1, starch synthase7; SS2c, starch synthase6; Sbe1, starch branching enzyme1; Bt1, brittle endosperm1; Ae1, amylose extender1; SS2b-2, starch synthase homolog1; Su2, sugary2; Isa2, sugary4; Gbss1, granule-bound starch synthase1; Isa3, isoamylase-type starch debranching enzyme3; Amyb5, beta amylase5; SS4, starch synthase4; Sbe3, starch branching enzyme3; SS1, Starch Synthase 1; Wx1, waxy1; Du1, dull endosperm1; SS3b-1, Starch Synthase 3; Sus2, sucrose synthase2; Pdh2, pyruvate dehydrogenase2; Mdh4, malate dehydrogenase4; Pgm1, phosphoglucomutase1; Phi1, phosphohexose isomerase1; Incw4, invertase cell wall4; Ivr1, invertase1; Mn1, miniature seed1; Pdh1, 6-phosphogluconate dehydrogenase-2; Ivr2, invertase2; Incw1, cell wall invertase1; Pgd1, 6-phosphogluconate dehydrogenase1; Mdh2, malate dehydrogenase2; Ra3, ramosa3; Tpi3, triose phosphate isomerase3; Spp1, sucrose-phosphatase1; ZmHxk3b, hexokinase9; Sps1, sucrose phosphate synthase1; Pmg1, phosphoglycerate mutase1; Sh1, shrunken1; Sus1, sucrose synthase1; ugp1, UDP-glucose pyrophosphorylase1; ugp2, UDP-glucose pyrophosphorylase 2; mips1, myo-Inositol-1-phosphate synthase 1; mips2, myo-Inositol-1-phosphate synthase 2; mips3, myo-Inositol-1-phosphate Synthase 3; gols1, galactinol synthase 1; gols3, galactinol synthase 3; rafs1, raffinose synthase 1; rafs2, raffinose synthase 2; rafs3, raffinose synthase 3; aga5, stachyose synthase; aga1, alkaline alpha-galactosidase 1; aga2, alkaline alpha-galactosidase 2; aga3, alkaline alpha-galactosidase 3.

GO enrichment analysis revealed that down-regulated genes in *wx-sweet* were significantly enriched in carbohydrate metabolism-related pathways (Supplementary Dataset S3), such as “pyruvate metabolic process” and “glycolytic process", particularly at 17 DAP. In contrast, up-regulated genes were significantly enriched in carbohydrate or lipid metabolic processes at multiple stages. Notably, at 14 DAP, up-regulated genes in *wx-sweet* were significantly enriched in several GO terms, including “carbohydrate metabolic process,” “carbohydrate catabolic process", “glycogen metabolic process", “monosaccharide metabolic process", and “glucan metabolic process” (Fig. 5C; Supplementary Dataset S3).

The decline in *Sh1* expression led to altered expression of sucrose metabolism-related genes across multiple developmental stages. The up-regulation of cell wall invertase genes *Mn1* (CWI-2) and *Incw1* suggests that the loss of *Sh1* function may induce *Mn1* expression through a sucrose accumulation-mediated feedback mechanism (Fig. 5F; Supplementary Fig S4; Supplementary Table S3). A key raffinose metabolism gene, Zm00001d021249 encoded a bifunctional UDP-glucose 4-epimerase and UDP-xylose 4-epimerase 1, was up-regulated 1.27-fold and 2.14-fold at 17 and 23 DAP, respectively, while *Rafs3* exhibited the greatest up-regulation (7.29-fold) at 26 DAP (Fig. 5H; Supplementary Table S3). Phosphogluconate dehydrogenase (Pdh1) and malate dehydrogenase 2 (Mdh4), key enzymes in the pentose phosphate pathway and tricarboxylic acid cycle, respectively, also showed up-regulated trends in *wx-sweet*. Furthermore, the expression of several known starch synthesis-related genes, including *Bt1*, *Sh2*, *Bt2*, *SS1*, and *Su1*, was altered (Fig. 5G; Supplementary Table S3). These results indicate that *Sh1* deficiency triggers a reprogramming of carbohydrate metabolism in the endosperm, potentially leading to substantial sugar accumulation and altered amylopectin synthesis in *wx-sweet*.

Notably, genes involved in the benzoxazinoid defense metabolism pathway, particularly those related to the synthesis of the strongly sweet-tasting compound DIMBOA-Glc, were generally up-regulated in *wx-sweet*. Among them, *Bx4*, *Bx5*, and *Bx6*, which collectively catalyze DIMBOA production, were significantly up-regulated during mid to late stages (20–26 DAP). Their modifying genes, *Bx10*, *Bx11*, *Bx13*, and *Bx14*, were also significantly up-regulated at 20 and 23 DAP (Fig. 5I; Supplementary Table S3), suggesting that this pathway may contribute to the sweet taste phenotype of the mutant.

### 3.6 Sugar Metabolomic Profiling Reveals Substantial Alterations in Sugar Metabolism in the *sh1* Endosperm

To elucidate the role of *sh1* in regulating sugar metabolism, we conducted targeted GC– MS-based metabolomic analysis to compare the sugar metabolic profiles between *wx* and *wx-sweet* kernels at 14, 20, and 23 DAP (Fig. 6; Supplementary Table S4). Among the 32 annotated sugar-related metabolites (Fig. 6A), 13 differentially accumulated metabolites (DAMs, criteria: |log₂FoldChange| > 0.5, FDR < 0.05) were identified at 17 DAP, 12 of which were significantly downregulated in *wx-sweet*. At 20 DAP, 11 DAMs were detected, with eight showing significant downregulation in *wx-sweet*. By 23 DAP, the number of DAMs in *wx-sweet* endosperm decreased to only four. Although the accumulation levels of multiple sugar metabolites were significantly reduced in *wx-sweet*, total sugar content was higher than in *wx*. This phenomenon can be attributed to the significantly higher sucrose content in *wx-sweet* across all three developmental stages compared to *wx*, although the log2 (Fold Change) < 0.5 (Fig. 6B; Supplementary Table S4). While, fructose and glucose levels were significantly lower in *wx-sweet* at all three time points (Fig. 6B). This metabolic profile likely results from reduced sucrose synthase activity due to the *sh1* mutation. In addition to increased sucrose, raffinose accumulation was also significantly elevated in *wx-sweet* kernels throughout all three developmental stages. This result is consistent with the upregulated expression of key raffinose metabolism genes identified in RNA-seq analysis, providing further metabolic evidence that loss of *Sh1* function regulates sugar accumulation.

**Figure 6.**
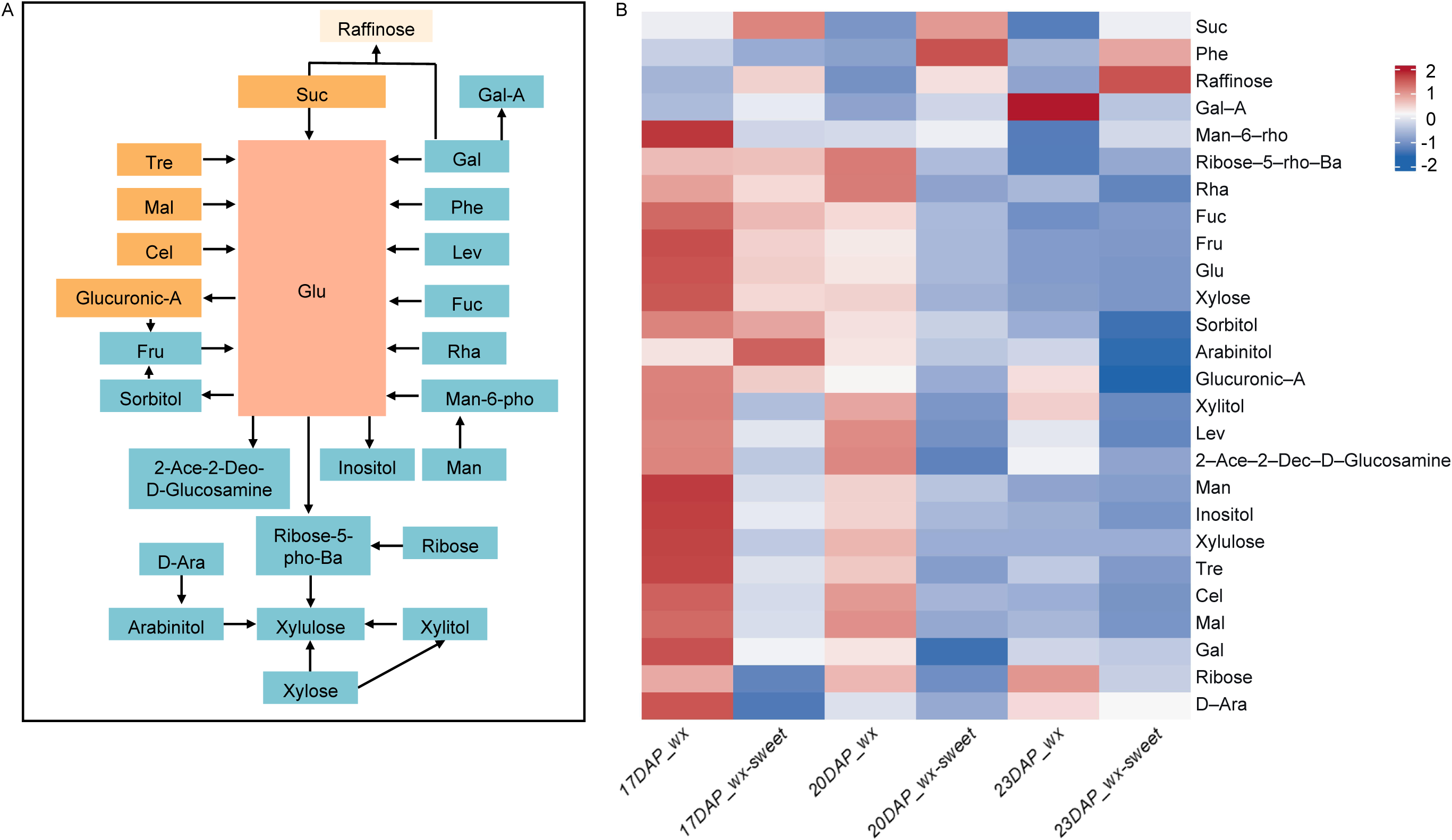
Sugar metabolomic profiling in the *wx-sweet* and *wx* endosperms. A. Pathway related to sugar metabolic pathway in maize. B. Heatmap of sugar metabolite content at different stages (17, 20, and 23 DAP) in *wx-sweet* and *wx* endosperms.

### 3.7 Application of the novel allele of *sh1* in Breeding Waxy-Sweet Corn

Based on the 82-bp insertion identified at the +415 bp site in *sh1* gene of the *wx-sweet* mutant (Fig. 3B; Supplementary Table S2), we developed an InDel marker (*Sh1*-InDel) spanning this insertion site (Supplementary Table S5). To validate this marker, we used it to genotype relevant materials. As expected, it amplified a 657-bp fragment in the *wx* seedlings, a 739-bp fragment in the *wx-sweet* seedlings, and both fragments in their F₁ (*wx* × *wx-sweet*) hybrid seedlings (Fig. 7A). We performed backcrossing using two elite waxy maize inbred lines (N-09, and N-29) as recurrent parents, and *wx-sweet* as the donor (Fig. 7B, D). The Sh1-InDel marker was used for genotypic selection of progeny plants carrying the novel allele of *sh1*. After five consecutive backcrosses followed by two generations of self-pollination, the inbred line 24H09 (Fig. 7C), homozygous for the *sh1* allele, was obtained in the N-09 background (Fig. 7E), and the inbred line 24H29 was obtained in the N-29 background. The hybrid F₁, named ‘Su nuo wei tian 218’, was produced by crossing 24H09 and 24H29 (Fig. 7G).

**Figure 7.**
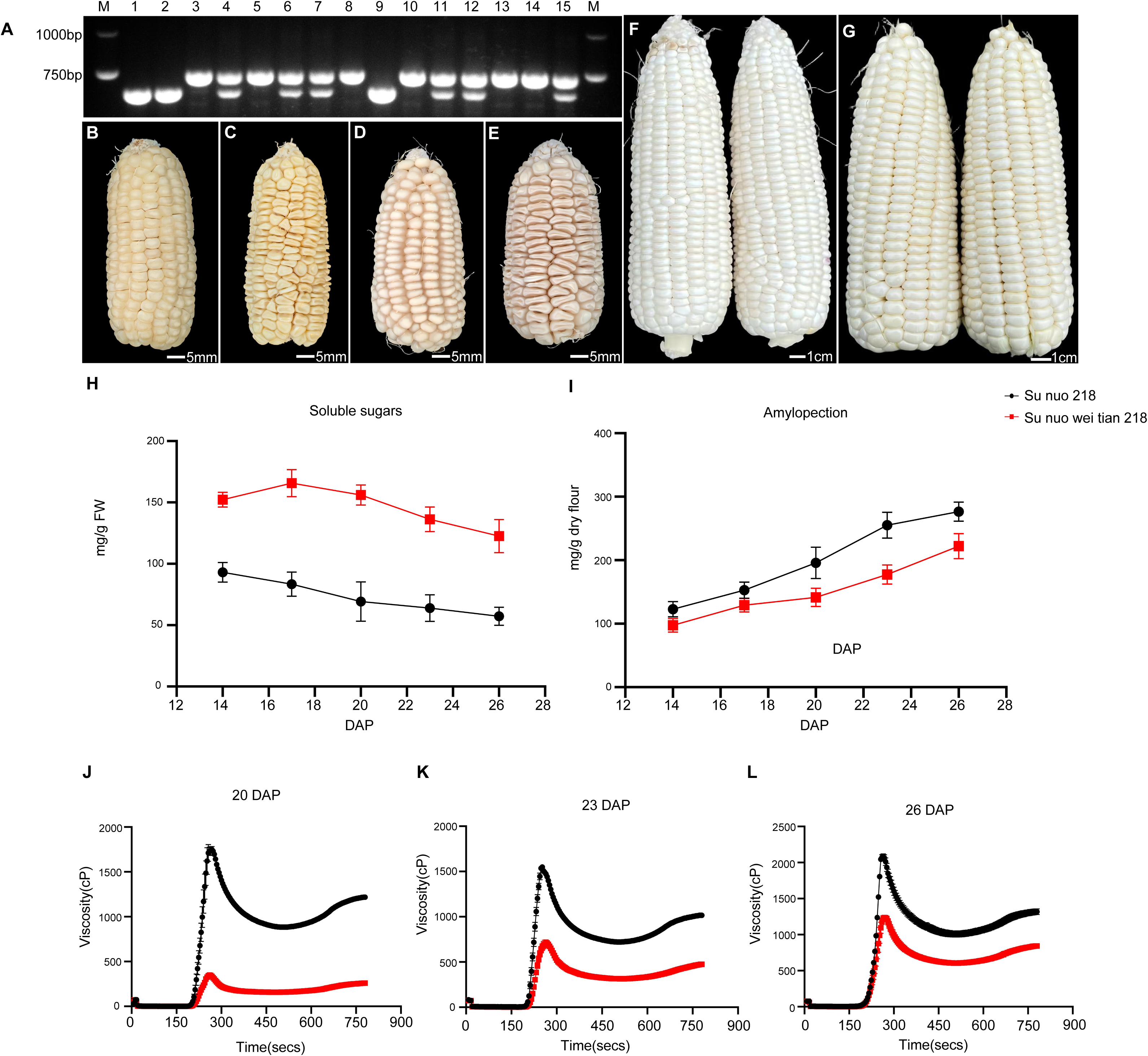
A Sweet-Waxy Varieties by Backcrossing with *sh1* in this study. A: Selection using functional molecular markers for *sh1* during backcrossing; B: The ear of N-29 (*waxy1waxy1*) at the mature stage, scale bar = 5mm; C: The ear of 24H29 (*waxy1waxy1*sh1sh1) at the mature stage, scale bar = 5mm; D: The ear of N-09 (*waxy1waxy1*) at the mature stage, scale bar = 5mm; E: The ear of 24H09 (*waxy1wax1sh1sh1*) at the mature stage, scale bar = 5mm; F: The ear of F_1_ (N-29 × N-09), ‘Su nuo 218’, at 23 DAP, scale bar = 10mm; G:The ear of F_1_ (24H29 × 24H09), ‘Su nuo wei tian 218’, at 23 DAP, scale bar = 10mm; H: Soluble sugar content in kernels of F_1_ (N-29 × N-09) and F_1_ (24H29 × 24H09) at different developmental stages; I: Amylopectin content in kernels of F_1_ (N-29 × N-09) and F_1_ (24H29 × 24H09) at different developmental stages; J–L: Pasting properties in kernels of F1 (N-29 × N-09) and F1 (24H29 × 24H09) at different developmental stages.

We then measured the soluble sugar content in the endosperm of ‘Su nuo wei tian 218’ at 14, 17, 20, 23, and 26 DAP (Fig. 7H). The content showed a trend of first increasing and then decreasing, remaining above 15% at all stages except 26 DAP, and peaking at 17 DAP (182.46 mg/g). In contrast, the soluble sugar content of the control hybrid ‘Su nuo 218’ (N-09 × N-29) was consistently lower than that of ’Su nuo wei tian 218’ during the same period and exhibited a continuous decline (Fig. 7H). Furthermore, we measured amylopectin content of ‘Su nuo 218’ and ‘Su nuo wei tian 218’. While it gradually accumulated in both ‘Su nuo 218’ and ‘Su nuo wei tian 218’ with the time after pollination, the latter showed significantly lower levels at all time points (Fig. 7I). To further compare their waxy properties, we examined the pasting properties of kernels at 20, 23, and 26 days after pollination (DAP). The results showed that the pasting curves of ‘Su nuo wei tian 218’ were consistently lower than those of ‘Su nuo 218’ at all time points (Fig. 7J-L). However, the difference in pasting curves between the two varieties gradually diminished during the developmental progression. These findings demonstrate that ‘Su nuo wei tian 218’ successfully combines sweetness and waxy texture during the harvesting period (20 – 26 DAP). Through this breeding strategy, we have developed a novel maize material possessing both sweet and waxy characteristics.

## 4 Discussion

The pursuit of high-quality sweet-waxy corn has gained significant momentum in breeding programs, driven by consumer demand for varieties that combine sweet taste with a waxy texture in a single kernel. In this study, we isolated a novel high-sugar waxy maize mutant (Fig. 1; Fig. 2), *wx-sweet*, through fast neutron mutagenesis and identified the causal mutation in the *Sh1* gene via map-based cloning (Fig. 3).

### 4.1 A Novel Allele of *sh1* Identified in the *wx-sweet* Mutant

The *Sh1* locus encodes sucrose synthase, which catalyzes the cleavage of sucrose into UDP-glucose and fructose. Numerous *sh1* mutant alleles have been reported from both natural populations and artificial mutagenesis, all impacting starch and sugar metabolism in maize kernels. For instance, EMS-induced alleles and the spontaneous mutant sh-7107 retain only about 10% of wild sucrose synthase activity (Chourey and Nelson, 1971). Other alleles, such as sh-bz-m4 with a Ds transposon insertion and sh9026 with a Mu insertion in the first exon, result in a complete loss of function or disruption of mRNA transcription and processing (Chourey and Nelson, 1976; Anderson et al., 1991; Ortiz et al., 1988). The natural allele *sh1-m* carries a 5-bp substitution causing a frameshift and premature termination, leading to a complete loss of enzyme activity and elevated soluble sugar content (Guan et al., 2017). Similarly, the EMS-induced *sh1* allele featuring a missense mutation (H433Y), results in a > 90% reduction in sucrose synthase activity, significantly reducing starch and increasing soluble sugar levels (Zhang et al., 2020). Two other EMS-induced mutants identified by Deng (2020) harbor G→A transitions at intron splice acceptor sites, causing aberrant RNA splicing. It is noteworthy that while sucrose synthase activity is drastically reduced in this mutants (Deng et al., 2020), the residual activity levels are not uniformly as low as the ∼10% previously reported. Our study has also observed similar result (Fig. 4A). It is possible that sucrose synthase activity is affected by currently undocumented sucrose synthase-encoding genes in maize, such as sus3 (Fig. 4A).

However, a critical research gap remained, as prior functional studies of *Sh1* were predominantly conducted in normal starch maize backgrounds. Its role and the phenotypic consequences of its loss in a *waxy1* genetic background, where starch metabolism is inherently altered, were unclear. Our work addresses this gap by reporting the first artificially induced *sh1* allele created directly within a waxy maize inbred line. This new allele confers a highly desirable dual phenotype: significantly enhanced sweetness while successfully retaining the essential waxy texture. This finding demonstrates the feasibility of engineering sweet-waxy traits and provides a valuable genetic resource for breeding next-generation fresh-eating corn varieties.

### 4.2 Complex Interplay Between Sweetness Genes and the *waxy1* gene

In earlier breeding attempts, researchers crossed common sweet corn genes (e.g., *su1*, *sh2*, *bt2*, *bt1*, and *du1*) with the *waxy1* gene to develop kernels combining both sweet taste and waxy texture. Although these sweet genes exert specific metabolic effects in the endosperm, their combinations with *waxy1* resulted in distinct epistatic interactions, ultimately shaping the final sugar and starch profiles. Studies confirmed that in homozygous double or multiple mutants, amylose content was significantly reduced (Creech, 1965; Wu, 2003), indicating that the waxy phenotype conferred by the *waxy1* gene was partially expressed. However, in sensory evaluations, these double mutants generally failed to exhibit both desirable sweetness and a waxy mouthfeel simultaneously.

For instance, the *sh2* gene encodes the large subunit of ADP-glucose pyrophosphorylase, a key rate-limiting enzyme in starch synthesis (Ballicora et al., 2004). The *sh2* mutation strongly inhibits starch biosynthesis, resulting in very low amylopectin content. Consequently, although soluble sugar content in *sh2waxy1* double homozygous mutants is higher than in the *sh2* single mutant, their amylopectin level is markedly lower, causing their eating quality to resemble that of *sh2* corn, essentially lacking waxy texture (Holder et al., 1974; Wang et al., 2019). Similarly, *bt1* and *bt2*, which also function upstream of *waxy1* in the starch synthesis pathway, exhibit epistasis over *waxy1*, leading to *bt1waxy1* or *bt2waxy1* double mutants that similarly lack a pronounced waxy mouthfeel (Li et al., 2017). The *Du1* gene encodes soluble starch synthase SSSII. When *du1* is combined with *waxy1*, the conversion of ADP-glucose into both amylopectin and amylose is blocked, resulting in elevated total sugar content but drastically reduced starch. Thus, *du1waxy1* kernels display sweetness but lack waxiness (Kramer et al., 1958). On the other hand, *Su1* encodes isoamylase, which acts downstream of the granule-bound starch synthase encoded by *waxy1* in the starch synthesis pathway. The su1wx double mutant exhibits significantly lower sugar content than the *su1* single mutant (Creech, 1965), making it also unsuitable as a sweet-waxy corn.

In summary, despite extensive efforts to combine various sweet genes with *waxy1*, no homozygous genotype has been obtained that delivers an ideal sweet and waxy sensory experience, and the successful development of such varieties through these conventional approaches remains unreported in practical breeding.

### 4.3 The Molecular Mechanism Underlying the Simultaneous Sweetness and Waxiness texture in *wx-sweet* kernels

To elucidate the mechanism underlying the dual sweet and waxy texture of *wx-sweet* kernels, we systematically analyzed sugar and starch metabolic pathways through transcriptomic profiling and sugar metabolomics profiling (Fig. 5; Fig 6) . Compared with *wx*, the *wx-sweet* mutant exhibited significantly downregulated *Sh1* expression and a marked reduction in sucrose synthase activity, indicating a blockage in the *Sh1*-mediated sucrose cleavage pathway. Measurements of soluble sugar content across developmental stages revealed that *wx-sweet* accumulated 1.17 to 1.63-fold more sugar than *wx* (Fig. 6B; Fig. 7 H; Supplementary Table S3), forming the primary basis for its sweetness. Second, raffinose, a non-reducing sugar that minimally participates in the Maillard reaction, has recently been implicated in the flavor profile of fresh corn (Luo et al., 2024). Our analysis showed that several genes in the raffinose metabolic pathway (Zm00001d021249, *Rafs3*, *Aga2*, *Aga3*) were consistently upregulated across all five developmental stages in *wx-sweet* (Fig. 5H; Supplementary Table S3) . This suggests that the metabolic remodeling of starch and sucrose metabolism induced by the *sh1* mutation further activates raffinose biosynthesis, potentially contributing to the sweet taste through a secondary pathway. Third, although not a sugar, the benzoxazinoid derivative DIMBOA exhibits a sweetness intensity approximately 400 times that of sucrose (Hamilton et al., 1962; Luo et al., 2024; Zhou et al., 2018). In this study, key genes in the DIMBOA biosynthesis pathway (*Bx4*, *Bx5*, *Bx6*, *Bx7*, *Bx8*) were generally upregulated in *wx-sweet* across developmental stages (Fig. 5I; Supplementary Table S3), indicating that accumulation of this metabolite provides a third contribution to the strong sweet perception.

Regarding the maintenance of the waxy texture, loss of Sh1 function led to reduced UDP-glucose supply (Chourey and Nelson, 1976), thereby limiting ADP-glucose (ADPG) synthesis. In response, two compensatory mechanisms were activated in the *wx-sweet* endosperm. On one hand, the cell wall invertase genes *Mn1* and *Incw1* were upregulated (Fig. 5F; Supplementary Fig. S4; Supplementary Table S3), enhancing sucrose hydrolysis to replenish hexose precursors (Cheng et al., 1999; Xu et al., 1996). On the other hand, ADPG synthesis-related genes (*Bt1*, *Sh2*, and *Bt2*) were consistently upregulated from 17 to 23 DAP (Fig. 5G; Supplementary Table S3), increasing ADPG production capacity at the transcriptional level (Zhang et al., 2020). Concurrently, significant upregulation of *SS1* and *Su1* in the amylopectin synthesis pathway during mid to late developmental stages supported sustained amylopectin production.

In conclusion, the combination of *sh1* and *waxy1* achieves an ideal balance between sweetness and waxy texture through multi-pathway coordination. While the *sh1* mutation promotes sucrose accumulation and enhances raffinose and DIMBOA synthesis, it also activates compensatory mechanisms involving substrate supply, ADPG synthesis, and amylopectin assembly to maintain sufficient amylopectin content and waxy texture. This genetic interaction model provides a theoretical foundation and key germplasm resources for targeted breeding of sweet-waxy corn.

### 4.4 Development and Application of a Functional Marker of *sh1* for *wx-sweet*

The InDel marker developed in this study enables accurate identification of heterozygous (*Sh1sh1*) plants at the seedling stage. In backcross progenies, heterozygous and dominant homozygous (*Sh1Sh1*) kernels are phenotypically indistinguishable from *wx* kernels, making phenotypic selection impractical. Our marker facilitates efficient selection, thereby significantly saving time and resources in breeding programs. Using this marker, we have successfully introgressed the *sh1* allele identified here into elite Chinese waxy maize backgrounds, ultimately leading to the development of a new sweet-waxy corn variety.

In summary, this work not only provides a valuable genetic resource for sweet-waxy maize breeders but also offers a functional molecular marker for precise genotyping of the *sh1* locus, which will accelerate the breeding process. Additionally, our findings enhance the mechanistic understanding of gene interactions within starch metabolic pathways in a waxy maize genetic background.

## Author Contributions

HZ, HXD, and LHN designed the experiment. LZ, SQL, QQH, WMZ and YPC collected the phenotypes and performed the data analysis. LZ, and YXP participated in fine-mapping. JH and YCW participated in RNA isolation and RNA-seq analysis. Metabolomic analysis was conducted by ZFG. Collections of all plant materials were performed by HXD and QQH. LHN wrote the initial manuscript. HZ and HXD revised this draft. All authors have read and approved the manuscript.

## Declarations

Conflict of interest The authors declare that they have no known competing financial interests or personal relationships that could have appeared to influence the work reported in this paper.

## Supplementary Data

The following materials are available in the online version of this article.

**Supplementary Figure S1**. Iodine-staining of starch and gene expression profiling of *waxy1* and *Sh1* in *wx* and *wx-sweet* kernels.

**Supplementary Figure S2**. PCR amplification of the *Sh1* gene using genomic DNA and cDNA from *wx-sweet*.

**Supplementary Figure S3**. GO enrichment of differentially expressed genes (DEGs) in *wx-sweet* vs *wx* kernels at five developmental stages.

**Supplementary Figure S4**. Heatmaps of log2FoldChanges for invertase-encoding genes in maize.

**Supplementary Table S1**. The functionally annotated gene identified in the target region.

**Supplementary Table S2**. The variations of *sh1* in *wx-sweet*. **Supplementary Table S3**. The genes used this study for heapmap. **Supplementary Table S4**. Sugar metabolites list by GC-MS. **Supplementary Table S5**. Primers used in this study.

**Supplementary Data Set S1**. RNA-seq analysis of *wx* and *wx-sweet* kernels at five developmental stages.

**Supplementary Data set S2**. The DEGs of *wx* and *wx-sweet* kernels at five developmental stages.

**Supplementary Data set S3**. Enriched GO terms for significant DEGs of *wx* and *wx-sweet* at five developmental stages.

**Appendix S1.** The gene sequence of *Sh1* in *wx* and *wx-sweet*.

