## Supplementary material for "A novel allele of *Sh1* underlies the conversion of *waxy* corn to *wx-sweet* corn": Data: Supplementary Figure..docx

Supplementary Data. Zhou et al. (2025). Theoretical and Applied Genetics

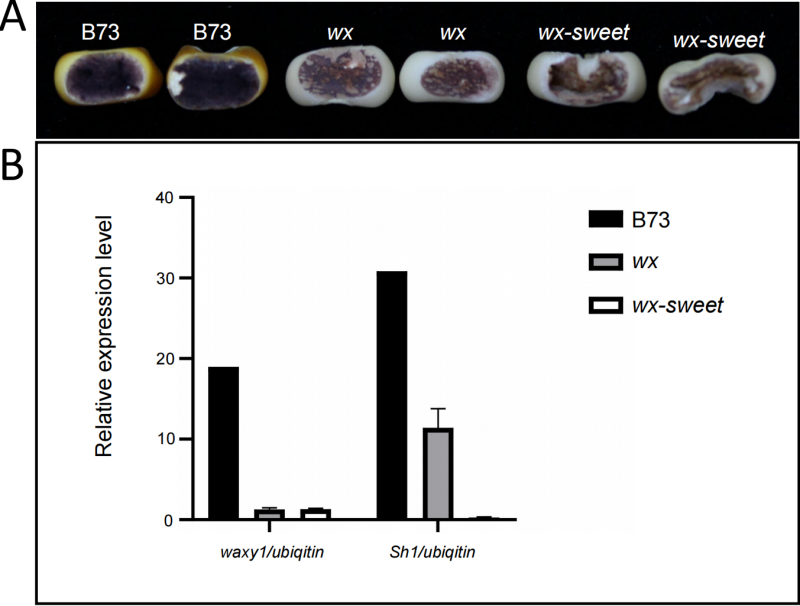

**Supplementary Figure S1.** Iodine-Staining of starch and gene expression profiling of *waxy1* and *Sh1* in *wx* and *wx-sweet* kernels.

A:Endosperm phenotypes of kernels following iodine-staining.

B: Expression levels of *Waxy1* and *Sh1* (presented as TPM ratio relative to Ubiquitin) in B73, *wx*, and *wx-sweet* kernels. TPM values for *wx* and *wx-sweet* were obtained from RNA-seq analysis of 14 DAP kernels in this study, while values for B73 were derived from published RNA-seq data of 14 DAP endosperm (Chen et al., 2014).

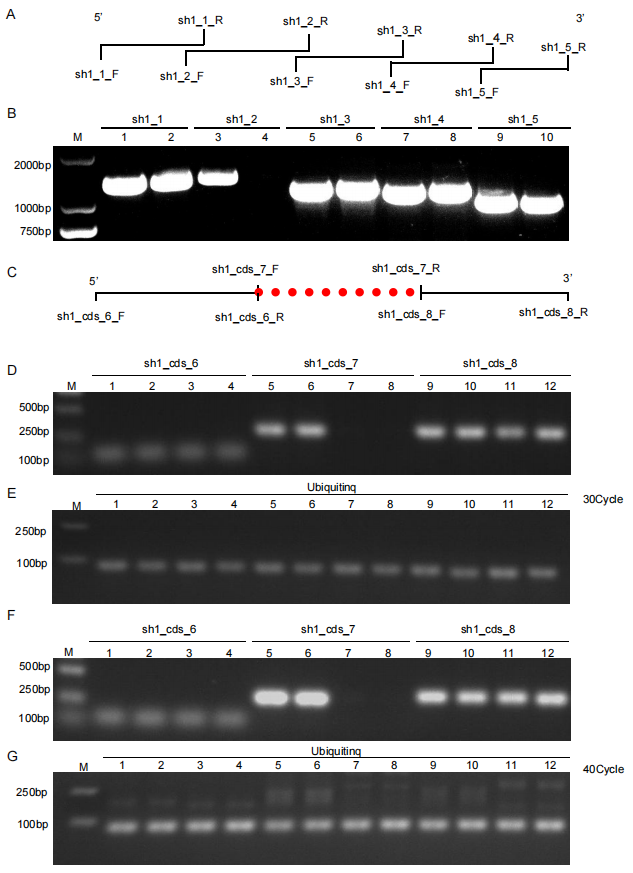

**Supplementary Figure S2.** PCR amplification of the *Sh1* gene using genomic DNA and cDNA from *wx*-*sweet* and *wx*.

A: Schematic representation of the *Sh1* gene structure and primer design. Five primer pairs (sh1_1_F/R to sh1_5_F/R) were designed based on the *Sh1* sequence of *wx* genome from second-generation sequencing to amplify the full-length gene. The diagram illustrates the regions covered by each primer pair and the expected amplification.

B: PCR amplification of DNA templates using the five primer pairs. Samples 1, 3, 5, 7, and 9 correspond to *wx* samples, while samples 2, 4, 6, 8, and 10 correspond to *wx-sweet* samples. The *wx-sweet* sample showed no amplification with the sh1_2 primers, whereas amplification was observed in *wx* samples, suggesting a potential structural variation in the *wx-sweet* genome.

C: Amplification of cDNA using three *Sh1* CDS-specific primer pairs (sh1_cds_6/7/8). Only sh1_cds_7 failed to produce a product (indicated by the red dot), implying a possible structural alteration in the corresponding region.

D: Comparison of amplification efficiency between *wx* and *wx-sweet* cDNAs using the three CDS primers under 30 PCR cycles. The sh1_cds_7 primers amplified the expected fragment from *wx* but not from *wx-sweet*, consistent with the result in C.

E: Amplification with Ubiquitin primers as an internal control. Successful amplification in all samples confirmed consistent RNA quality and cDNA synthesis efficiency, ruling out technical artifacts.

F, G: The same amplifications targeting the *Sh1* CDS region and the Ubiquitin control as shown in D and E, performed with 40 PCR cycles. D-G: Samples 1, 2, 5, 6, 9, and 10 are *wx*; samples 3, 4, 7, 8, 11, and 12 are *wx-sweet*. Increased cycle numbers resulted in clearer bands, reinforcing the above conclusions.

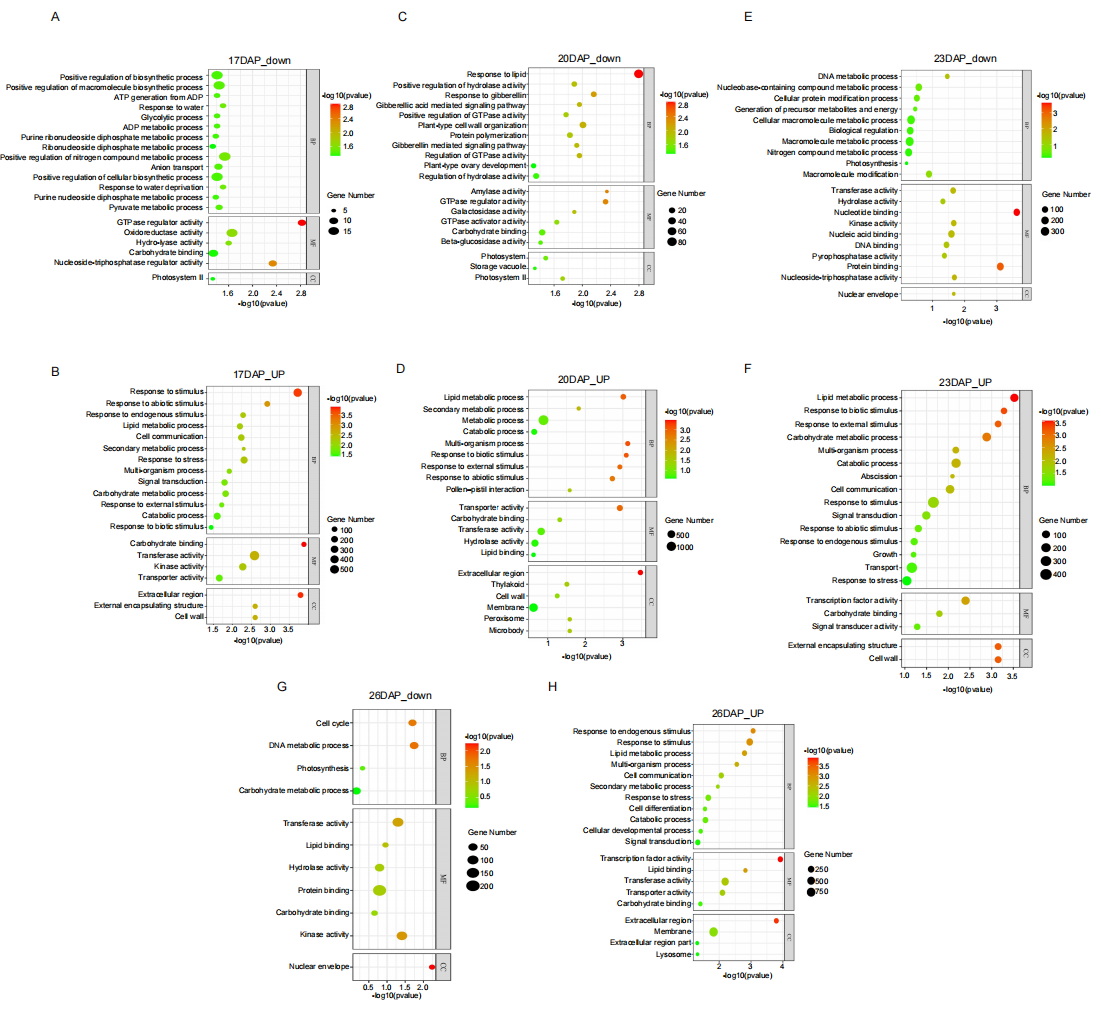

**Supplementary Figure S3.** GO enrichment of differentially expressed genes (DEGs) in *wx-sweet* vs *wx* kernels at five developmental stages (17, 20, 23, and 26 DAP). Y-axis label colors denote the significantly enriched GO terms. The color and size of the plotted symbols correspond to the enrichment significance level and the number of associated DEGs, respectively.

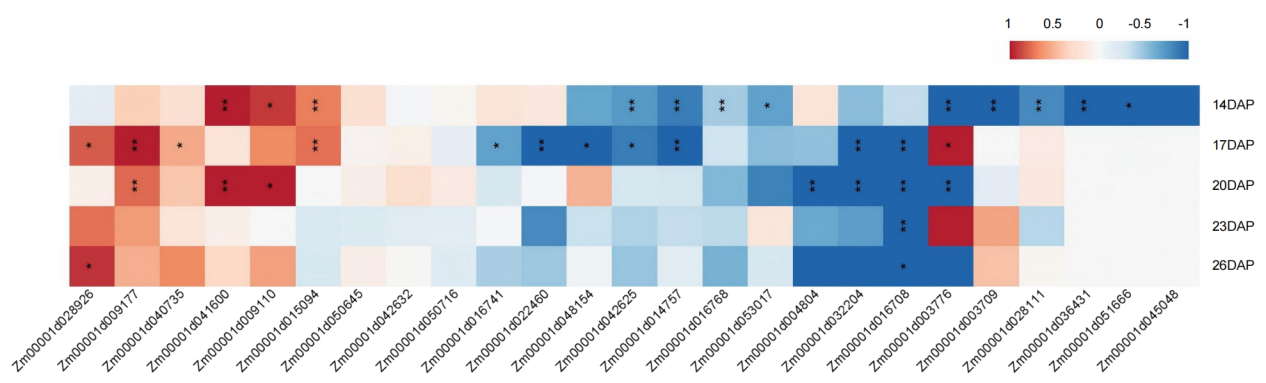

**Supplementary Figure S4.** Heatmaps of log₂FoldChanges for invertase-encoding genes in maize, as reported by Yi (2021). The heat maps display values (*wx-sweet* vs. *wx*) from 14, 17, 20, 23, and 26 DAP, which were used for clustering. See Supplementary Dataset 1 for further details.
