## Supplementary figures and images for "A novel allele of *Sh1* underlies the conversion of *waxy* corn to *wx-sweet* corn"

### Supplementary Figure S1. Iodine-staining of starch and gene expression profiling of waxy1 and Sh1 in wx and wx-sweet kernels..pdf

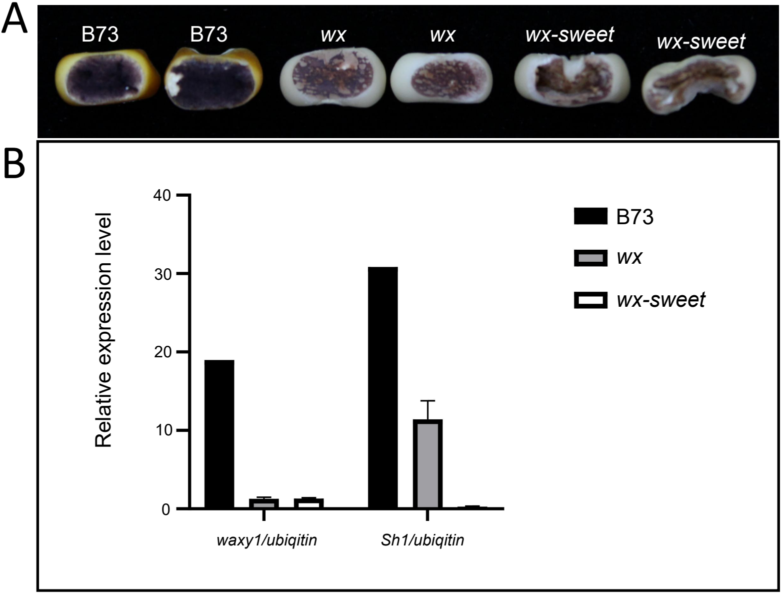

### Supplementary Figure S2. PCR amplification of the Sh1 gene using genomic DNA and cDNA from wx-sweet and wx..pdf

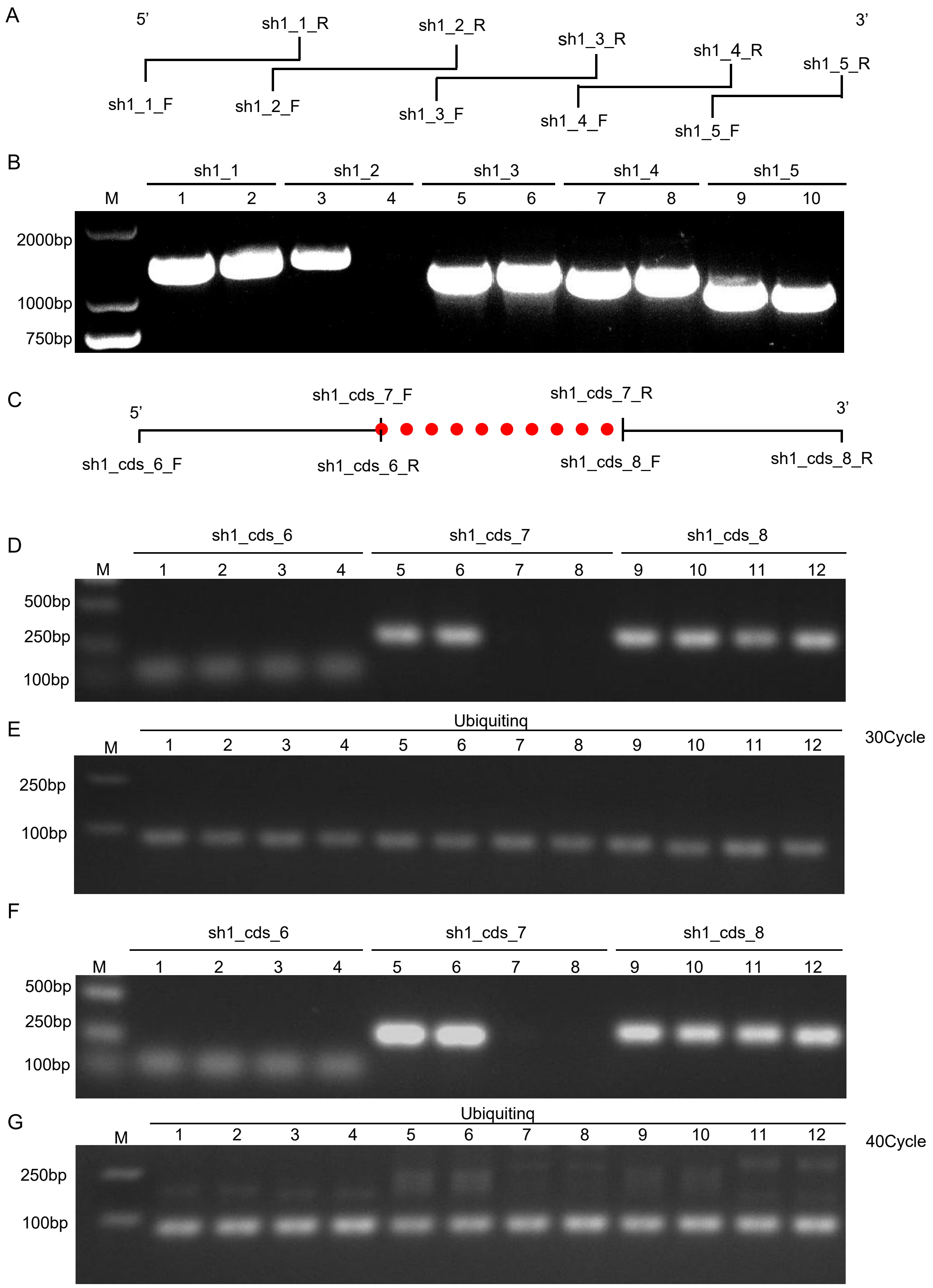

### Supplementary Figure S3. GO enrichment of differentially expressed genes (DEGs) in wx-sweet vs wx kernels at five developmental stages.pdf

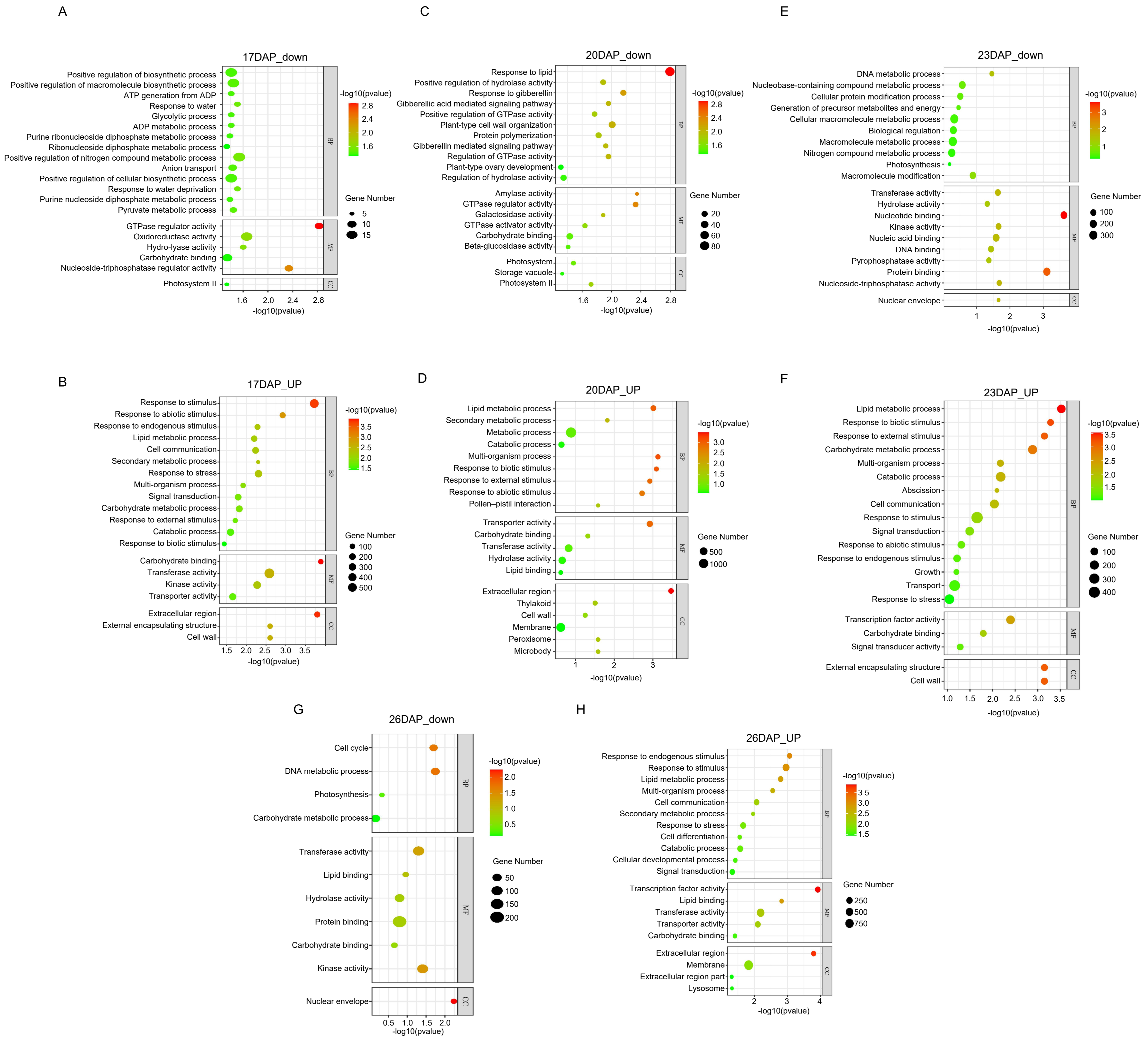

### Supplementary Figure S4. Heatmaps of log2FoldChanges for invertase-encoding genes in maize..pdf

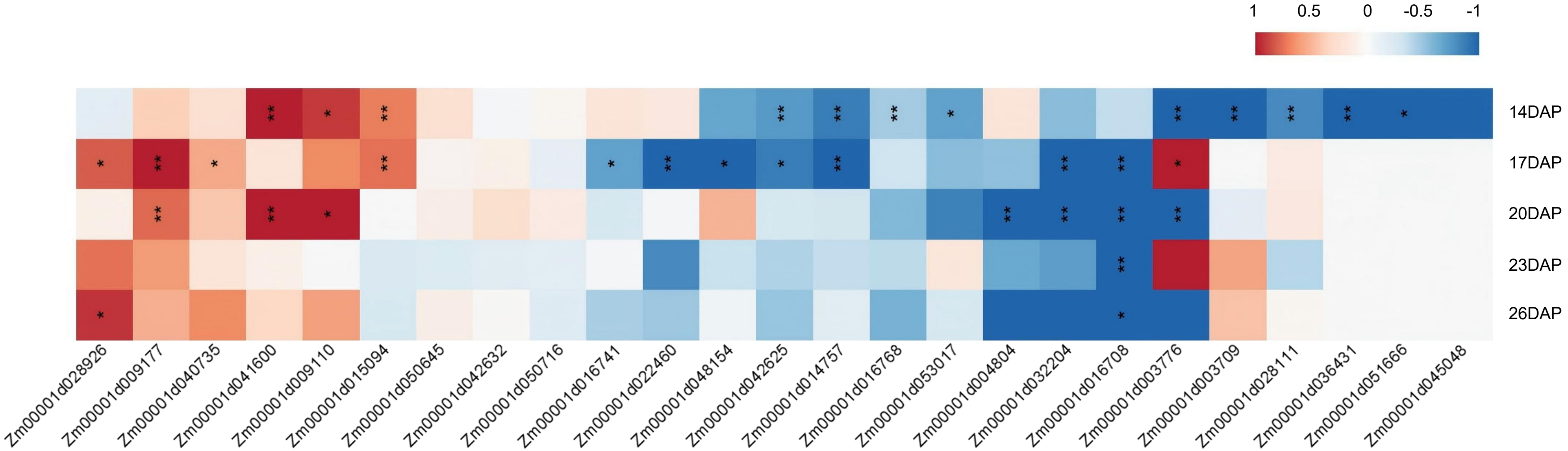
